# Molecular, cellular and network mapping of brain structural deviations in patients with Post-COVID19 syndrome

**DOI:** 10.64898/2026.01.12.699045

**Authors:** Daniel Martins, Ziyuan Cai, Nicole Mariani, Alessandra Borsinni, Valeria Mondelli, Brandi Eiff, Silvia Rota, Daniel van Wamelen, Timothy Nicholson, Laila Raida, Adam Hampshire, Lia Fernandes, Fernando Zelaya, Aleksandra Podlewska, Ray Chaudhuri, Federico Turkheimer, Steven C.R. Williams, Mattia Veronese

## Abstract

Post-COVID-19 syndrome encompasses persistent cognitive, neurological, and psychiatric symptoms following SARS-CoV-2 infection, profoundly affecting global quality of life. Clarifying the neurobiological basis of these symptoms is vital for effective therapeutic interventions. This study utilized normative modelling of brain structure (“CentileBrain”) to quantify individualized deviations in cortical thickness, surface area, and subcortical volumes among 20 patients experiencing persistent fatigue following mild COVID-19, compared to 20 matched healthy controls. Group-level analyses on deviation scores revealed subtle yet distinct regional alterations in cortical thickness, specifically decreased thickness within orbitofrontal cortices and increased thickness in occipital/sensory cortices. Although at the individual regional level, the proportion of patients exhibiting infranormal or supranormal thickness values was relatively low (<35%) and comparable to controls, deviations frequently clustered within structurally connected circuits, affecting up to 50% more of patients. Spatial analysis of regional cortical thickness alterations correlated significantly with the constitutive expression patterns of TMPRSS2, an essential protein facilitating SARS-CoV-2 cellular entry. Canonical correlation analyses further identified specific cell-type distributions and neuroreceptor densities predictive of regional thickness changes, highlighting neurons and molecular targets associated with serotoninergic, cannabinoid, cholinergic, and glutamatergic signalling pathways. Network-diffusion modelling constrained by a canonical structural connectome significantly outperformed null models based on permuted connectomes and Euclidean distance metrics, identifying posterior-parietal regions as probable initiation points (“seeds”) for network-wide structural changes. Seed likelihood correlated positively with TMPRSS2 expression levels, suggesting that these posterior-parietal regions may be particularly susceptible to SARS-CoV-2 infection. This highlights a plausible mechanism where structural alterations could propagate through connected neural networks, although direct evidence of such propagation requires further investigation. These findings provide novel insights into potential mechanisms underlying neural circuit disruptions in post-COVID-19 fatigue and suggest avenues for therapeutic neuromodulation.

**Highlights:**

1. Normative modelling revealed heterogenous individualised cortical structural deviations in patients with PCS.
2. Regional deviations in cortical thickness and surface area clustered within anatomically connected circuits, affecting up to 50% more of PCS participants.
3. Structural changes were significantly associated with regional TMPRSS2 expression, implicating SARS-CoV-2 entry pathways in PCS-related brain alterations.
4. Network diffusion modelling identified posterior-temporoparietal regions as likely epicentres of pathology propagation through the structural connectome.
5. Cortical deviations were linked to neuronal and microglial cell-type densities and receptor systems including serotonergic, glutamatergic, cannabinoid and cholinergic pathways, suggesting targets for neuroimmune-informed interventions.

## Introduction

Post-COVID-19 syndrome has rapidly emerged as a significant global health concern, characterized by an array of persistent cognitive, neurological, and psychiatric symptoms that occur following infection with SARS-CoV-2^1, 2^. Among these lingering symptoms, fatigue is notably prevalent and severely debilitating, adversely affecting daily functioning, productivity, and overall quality of life for millions worldwide^1–3^. Despite extensive clinical research and public health awareness, the precise neurobiological mechanisms underlying these persistent symptoms, particularly fatigue, remain poorly understood. This gap in knowledge critically impedes the development of targeted therapeutic interventions aimed at alleviating the burden of post-COVID-19 syndrome^2, 4, 5^.

Emerging evidence from neuroimaging studies has suggested that structural brain alterations might accompany even mild cases of COVID-19 infection. These alterations include subtle but potentially significant changes in cortical thickness, surface area, and subcortical brain volumes within regions involving olfactory, frontoparietal or limbic circuits^6–8^. However, existing findings are often inconsistent across studies, potentially due to methodological variability, small sample sizes, or the heterogeneity of patient populations. Moreover, traditional approaches typically rely on group-level analyses, which can obscure subtle individual differences in brain structure that may be clinically relevant. Consequently, a more nuanced approach capable of capturing individualized patterns of neuroanatomical deviation is necessary to better understand the underlying pathophysiological mechanisms^9–12^.

To address these limitations, our study employed a sophisticated normative modelling technique known as CentileBrain^9^. This method leverages large, demographically diverse reference datasets - including tens of thousands of healthy individuals - spanning a wide age range to establish robust, population-informed models of brain structure. Using Gaussian process regression, CentileBrain generates centile-based predictions of cortical and subcortical anatomy for each individual, accounting for covariates such as age, sex, and intracranial volume. By comparing observed brain morphometry to these normative expectations, the approach yields individualized deviation scores (z-scores), allowing detection of subtle structural abnormalities that may be clinically relevant but diluted in group-level comparisons^12, 13^. In this study, we applied CentileBrain to quantify deviations in cortical thickness, surface area, and subcortical volumes among patients with persistent fatigue following mild COVID-19. This personalized framework would allow us the identification of heterogeneously distributed and circuit-convergent brain alterations, offering deeper insight into the neuroanatomical basis of post-COVID syndrome.

Given that structural brain alterations alone do not directly reveal the underlying biological mechanisms, we sought to contextualize our neuroanatomical findings within molecular, cellular, and network-level frameworks^14, 15^. To this end, we tested a series of interrelated hypotheses designed to illuminate potential drivers of the structural changes observed in PCS. First, we examined whether regional deviations in cortical thickness and surface area were spatially correlated with the expression of genes implicated in viral entry and neuroimmune signalling, such as TMPRSS2, FURIN, and NRP1^16, 17^, which are central to viral entry mechanisms, and TLR4, a key component of the innate immune system involved in pro-inflammatory signalling triggered by the viral spike protein^18^. This analysis was intended to explore whether direct viral neurotropism or associated entry mechanisms contribute to regional structural vulnerability.

Second, we investigated whether these anatomical changes were associated with the spatial distribution of distinct brain cell types and neurotransmitter receptor systems, drawing on normative transcriptomic and PET-based datasets^14, 15^. We focused in particular on systems previously implicated in fatigue and neuropsychiatric dysfunction, including serotonergic, cannabinoid, cholinergic, and glutamatergic pathways. This approach aimed to identify whether specific neural populations or signalling systems are preferentially involved in PCS, thereby suggesting possible neurochemical mechanisms or therapeutic targets for prioritisation. Finally, we applied network-diffusion modelling using canonical structural connectivity data from the Human Connectome Project^19–23^. This allowed us to test whether focal structural deviations might propagate along anatomical pathways, giving rise to the distributed and heterogeneous patterns of brain alterations seen in PCS. By identifying putative epicenters and characterizing their diffusion dynamics, this approach sought to offers insight into how localized pathology may result in widespread circuit dysfunction and the multifaceted clinical symptoms characteristic of post-COVID fatigue.

## Methods

### Study Design and Participants

As described elsewhere, this study employed a single-site observational case-control design. All participants, both PCS cases and recovered controls, were required to have had biologically confirmed SARS-CoV-2 infection at least three months prior to enrolment, consistent with the WHO clinical case definition of Post-COVID-19 condition^24^. Individuals in the PCS group were characterised by persistent symptoms lasting ≥3 months after infection, whereas recovered controls were required to be fully asymptomatic, with complete resolution of all post-COVID symptoms. We included individuals with mild acute COVID-19 only, to minimise potential confounding effects of hospitalisation, hypoxia, or severe systemic inflammation on brain perfusion and metabolism, and to ensure that any observed group differences were not attributable to the neurological sequelae of moderate–severe disease. All participants were required to have had biologically confirmed SARS-CoV-2 infection prior to enrolment. Confirmation was based on either a positive reverse-transcription polymerase chain reaction (RT-PCR) test from a nasopharyngeal swab or documented serological evidence of prior infection, in accordance with UK national testing guidelines at the time of acute illness. Participants with only clinically suspected but biologically unconfirmed COVID-19 were not included. Recovered healthy control participants were required to have a history of biologically confirmed SARS-CoV-2 infection, followed by complete resolution of all acute and post-acute symptoms. To minimize the risk of including individuals with subtle or subclinical post-COVID manifestations, control participants were screened using the same standardized clinical instruments applied to the PCS group, including the Fatigue Assessment Inventory. Only individuals scoring below established clinical thresholds, and reporting no persistent fatigue, cognitive complaints, affective symptoms, autonomic dysfunction, or post-exertional malaise at the time of assessment, were included as healthy controls. In addition, control participants underwent the same cognitive assessment battery and clinical interview as PCS participants, and individuals demonstrating clinically relevant deviations from normative performance or reporting residual symptoms were excluded. This stringent screening strategy ensured that the control group represented a recovered post-COVID population without ongoing symptomatology, rather than asymptomatic or mildly affected PCS cases.

We recruited participants through King’s College Hospital NHS Trust and surrounding community networks. Eligible individuals were aged 16–65 years, fluent in English, and capable of providing informed consent. We excluded individuals with a history of major neurological or psychiatric disorders, current substance misuse, systemic or central nervous system inflammatory conditions, chronic respiratory or cardiac disease, recent vaccination (within two weeks), pregnancy or breastfeeding, BMI >30, or current use of immunomodulatory or psychoactive medications. We used the global severity score of Fatigue Assessment Inventory (FAI) to pre-screen participants for symptom severity ^25^. Individuals scoring greater than 4 on the global fatigue severity scale were classified into the PCS group with persistent fatigue, while those scoring 4 or below were included as recovered controls. The threshold of > 4 was selected to ensure inclusion of individuals with clinically meaningful levels of fatigue, consistent with prior applications of the FAI in post-viral and chronic fatigue contexts^25^. Groups were balanced for age, sex, BMI, and acute COVID-19 severity. The study received ethical approval from the UK Health Research Authority (IRAS ID: 308661) and was conducted in accordance with the Declaration of Helsinki and Good Clinical Practice (GCP) guidelines. All participants provided written informed consent prior to study procedures.

We conducted an a priori power analysis using G*Power (version 3.1) to estimate the required sample size for detecting group differences in dynamic functional connectivity metrics using analysis of covariance (ANCOVA). Assuming a medium to large effect size (Cohen’s f=0.30, equivalent to ηp^2^≈0.08), alpha = 0.05, power (1 − β) = 0.80, and three covariates (age, sex, and handedness), the estimated sample size required per group was 19. Our sample of 20 participants per group therefore provides adequate power to detect effects in this range.

### Procedures

Each participant completed a single study visit that included clinical assessments, venous blood collection, and a multimodal MRI scan. Clinical assessments included medical and psychiatric history, physical examination, vital signs, and a structured battery of standardized questionnaires. Fatigue was assessed using both the full Fatigue Assessment Inventory^25^ and the Multidimensional Fatigue Inventory^26^ to differentiate between mental and physical fatigue. Symptoms of autonomic dysfunction were evaluated using the Composite Autonomic Symptom Score (COMPASS-31)^27^, while mood and anxiety were measured using the Hospital Anxiety and Depression Scale (HADS)^28^. Sleep quality was assessed with the Pittsburgh Sleep Quality Index^29^, and respiratory symptoms were rated using the MRC Dyspnoea Scale^30^. Additional measures included the DePaul Symptom Questionnaire^31^, the American College of Rheumatology (ACR) fibromyalgia criteria^32^, the PAIN Detect questionnaire^33^, and Montreal Cognitive Assessment^34^. Most self-report questionnaires were completed remotely via the Qualtrics platform within 72 hours of the in-person visit. All participants underwent a structured diagnostic assessment using the Mini International Neuropsychiatric Interview (MINI) to exclude current or past major psychiatric disorders, including mood, anxiety, psychotic, and substance use disorders.

### Blood analysis

Venous blood samples (up to 30 mL) were collected on the day of imaging. Standard hematological and biochemical panels - including erythrocyte sedimentation rate (ESR), C-reactive protein (CRP), thyroid-stimulating hormone (TSH), free thyroxine (T4), and liver function tests - were processed by Synnovis (Viapath) at King’s College Hospital NHS Foundation Trust. Serum samples were frozen at −80°C and later assayed for cytokines and glial markers using electrochemiluminescence immunoassays (Meso Scale Discovery, Rockville, MD, USA). The cytokine panel included interleukin-1 beta (IL-1β), interleukin-2 (IL-2), interleukin-4 (IL-4), interleukin-6 (IL-6), interleukin-8 (IL-8), interleukin-10 (IL-10), interleukin-12p70 (IL-12p70), interleukin-13 (IL-13), tumor necrosis factor-alpha (TNF-α), and interferon-gamma (IFN-γ). Glial markers included glial fibrillary acidic protein (GFAP), measured with the ultrasensitive kit, and S100β. Assays were performed in duplicate. Values below the detection limit were discarded. This led us to discard all quantifications of IL-2, IL-4 and IL-12p70. Outliers greater than three standard deviations from the group mean were winsorized. No analyte had more than 10% missing data, therefore all data were included.

### Cognitron

Cognitive performance was assessed online using a battery of computerized tasks from the Cognitron platform, covering delayed verbal memory, working memory, executive function, attention, motor coordination, and spatial manipulation^35^. Tasks included Verbal Analogies (abstract reasoning and semantic integration), Immediate and Delayed Prospective Memory (episodic memory and planning), 2D Spatial Manipulation (visuospatial working memory and mental rotation), Motor Control (sensorimotor coordination and reaction consistency), Spotter (vigilance and sustained attention), and Block Reasoning (fluid intelligence and rule-based problem solving). Each task involves trial-level response collection, capturing both accuracy and reaction time. Performance metrics were transformed into standardized deviation-from-expected (DFE) scores using normative models derived from a large independent sample (n > 20,000), adjusted for age, sex, handedness, and harmonized ethnicity. DFE scores represent standardized residuals (observed minus expected performance divided by normative standard deviation) and were computed separately for accuracy and reaction time.

### Neuroimaging Acquisition and Processing

Magnetic resonance imaging (MRI) data were acquired using standardized imaging protocols on a 3T MRI scanner equipped with a standard head coil. High-resolution structural images were collected, including T1-weighted anatomical scans obtained using an MPRAGE sequence (isotropic voxel size = 1 mm³, TR = 2300 ms, TE = 2.98 ms, flip angle = 9°, field-of-view [FOV] = 256 mm) and complementary T2-weighted images (voxel size = 1 mm³, TR = 3200 ms, TE = 408 ms, flip angle = 120°, FOV = 256 mm). These sequences allowed detailed assessment of cortical thickness, cortical surface area, and subcortical brain volumes, leveraging the complementary tissue contrast provided by both imaging modalities. Image preprocessing and morphometric analyses were conducted using validated, automated pipelines implemented in FreeSurfer (version 7.2)^36^. T1-weighted images were initially processed through skull stripping to remove non-brain tissue, followed by bias field correction to mitigate intensity inhomogeneities. Subsequently, images underwent segmentation into gray matter, white matter, and cerebrospinal fluid. The complementary T2-weighted images were integrated into the FreeSurfer pipeline, enhancing the accuracy of pial surface delineation and improving segmentation quality, particularly in cortical regions and areas adjacent to cerebrospinal fluid.

Rigorous quality control (QC) procedures were implemented at multiple stages of the neuroimaging pipeline to ensure the accuracy and reliability of morphometric estimates. In addition to the default FreeSurfer QC procedures, we utilized the open-source FSQC tool developed by the Deep-MI initiative (https://deep-mi.org/fsqc/dev/index.html) to automate the detection of potential segmentation errors and anatomical outliers across multiple FreeSurfer outputs^37^. Each participant’s imaging data and segmentation outputs were subsequently subjected to detailed manual visual inspection by an experienced rater, using the FSQC interface alongside FreeSurfer’s built-in visualizations. Key features examined included motion artifacts, intensity non-uniformities, accuracy of white and pial surface placement, and the fidelity of cortical and subcortical parcellation. Datasets failing initial QC underwent manual correction, including adjustments to brain masks, removal of skull-strip artifacts, and refinements of white matter and pial boundaries. Corrected images were reprocessed through the FreeSurfer pipeline and re-evaluated to confirm improved segmentation quality. Only datasets that passed both automated and manual QC thresholds were retained for subsequent statistical analyses. This stringent multi-step approach ensured high-quality morphometric data and minimized the inclusion of artifacts or segmentation errors that could confound downstream individual-level deviation estimates or group-level comparisons.

### Normative Modelling Approach

To quantify individual deviations in brain structure, we employed the *CentileBrain* normative modelling framework, which estimates expected neuroanatomical features - cortical thickness, surface area, and subcortical volumes - based on a large reference dataset of healthy controls using Gaussian Process Regression^9^. Individual deviation scores were expressed as z-scores, reflecting how much a given regional brain measurement deviated from the normative prediction for that person’s age, sex, and intracranial volume. We defined extreme deviations as z-scores falling outside the ±1.96 threshold, corresponding to values in the top or bottom 2.5% of the normative distribution (i.e., below the 5th percentile or above the 95th percentile). For each cortical or subcortical region, we determined whether a participant exhibited an extreme deviation. This binarized information was used to calculate regional overlap, defined as the proportion of participants in each group (PCS or controls) showing extreme deviations in each region. Group differences in overlap were visualized as difference maps, where positive values indicated greater overlap in PCS and negative values indicated greater overlap in controls.

To examine whether these deviations converged within large-scale brain systems, we performed circuit-level overlap analyses. Following the principles of lesion-network mapping^38^, for each individual, we also identified regions with extreme deviations and then mapped their structural connectivity profiles using a normative structural connectome derived from diffusion-weighted imaging data in the Human Connectome Project^39^. For each deviated region, we extracted all directly connected regions (first-order neighbours) and combined them into a participant-specific circuit^40^. We then calculated the percentage of participants in each group who showed at least one extreme deviation within each circuit. Circuit-level overlap thus captured the frequency with which deviations occurred in anatomically connected subnetworks, even if the specific deviated nodes varied across individuals. Together, the regional and circuit-level analyses enabled us to assess whether the anatomical deviations observed in PCS were spatially random or showed consistent patterns of convergence within specific areas or networks. This multiscale framework allowed us to identify not only which brain regions were most frequently affected across individuals, but also whether these deviations reflected disruption of shared neural circuits^40^.

### Network-Diffusion Modelling

To examine the potential propagation pathways of structural brain alterations across neural circuits, we implemented network-diffusion modelling using structural connectivity data from the Human Connectome Project^19–23^. The diffusion modelling was based on a heat-kernel diffusion process applied to a structural connectome derived from diffusion-weighted imaging as previously reported^41^, repeating the seeding process iteratively from each region of the parcellation used at a time. The diffusion model involved optimizing the diffusion coefficient (β), which controls the rate of propagation within the connectome. Optimization of β was achieved by systematically evaluating diffusion performance across a range of values (0.1–1.0, increments of 0.1), selecting the β parameter that maximized the correlation between the simulated diffusion pattern and empirical neuroanatomical thickness deviations as it was in this metric where we found most differences. Diffusion simulations were constrained to reflect propagation over a biologically plausible time range (0–24 months post-infection), enabling investigation of both short-term and long-term diffusion dynamics. Permutation testing was performed to assess the statistical significance of the observed diffusion patterns. Specifically, empirical diffusion results were compared against null distributions generated by running diffusion simulations on scrambled connectomes with randomized connectivity patterns and Euclidean distance-based surrogate connectomes. A total of 1,000 permutations were conducted for each approach, establishing the statistical robustness of observed propagation patterns and identifying network hubs and pathways preferentially involved in structural alterations associated with prolonged fatigue following mild COVID-19 infection.

### Molecular and Cellular Analyses

Regional microarray expression data: Gene expression data from the AHBA were obtained from six adult postmortem brains aged 24–57 years. Genetic probes were reannotated following standardized methods outlined in *Arnatkeviciute et al. (2019)*^42^ to increase accuracy, and probes deemed unreliable were excluded. For each gene, the most stable probe was selected based on pooled donor correlations, resulting in a final dataset of 15,633 probes across the brain. Microarray samples were spatially assigned to the 83 regions of the Desikan-Killiany atlas40 using the abagen toolbox (https://abagen.readthedocs.io/en/stable/), with regional assignment based on corrected MNI coordinates^43^. Assignments were constrained by hemisphere and cortical/subcortical boundaries to improve anatomical accuracy. Gene expression values were normalized using a robust sigmoid function and rescaled to a unit interval for cross-region and cross-donor comparability. Normalization was performed separately for cortical and subcortical regions to circumvent know differences in mean gene expression between cortex and subcortex. Due to limited availability of samples from the right hemisphere, we mirrored samples from the left hemisphere into the right hemisphere to ensure all regions are assigned values of gene expression for all genes.

### Candidate gene analyses

To explore the molecular vulnerability of specific brain regions to SARS-CoV-2-related pathology, we examined whether regional variations in cortical thickness and surface area—derived from normative modelling - corresponded spatially with the constitutive expression of genes implicated in viral entry and neuroimmune signalling, as previously mapped using the same anatomical parcellation. We focused on a set of a priori candidate genes with well-established roles in mediating SARS-CoV-2 infection and its downstream inflammatory responses. TMPRSS2 (transmembrane serine protease 2) plays a critical role in priming the viral spike (S) protein, facilitating fusion of the viral and host cell membranes and enabling entry into host cells. TMPRSS2 is expressed in both respiratory and neural tissues and is thought to modulate region-specific susceptibility to direct viral invasion^44^. FURIN encodes a proprotein convertase that cleaves and activates the spike protein at the S1/S2 site, a step essential for viral infectivity. Its widespread expression in the central nervous system may contribute to facilitating viral spread in brain tissue, particularly in regions with high baseline FURIN activity^45^. NRP1 (neuropilin-1) has been identified as a co-receptor that enhances SARS-CoV-2 cell entry by stabilizing spike protein interactions with ACE2. Beyond its role in viral infectivity, NRP1 is also involved in axonal guidance and neurovascular signaling, raising the possibility that its expression could influence both entry pathways and downstream neurovascular dysfunction^16^. TLR4 (toll-like receptor 4) is a key component of the innate immune system that recognizes pathogen-associated molecular patterns (PAMPs) and initiates inflammatory cascades. Recent studies have shown that TLR4 may directly respond to the SARS-CoV-2 spike protein, leading to exaggerated pro-inflammatory responses. In the CNS, TLR4 activation has been implicated in microglial priming and neuroinflammation, which are candidate mechanisms for post-viral neuropsychiatric sequelae^46^. Together, these genes represent plausible molecular entry points and immunological amplifiers of SARS-CoV-2-related effects on the brain. Unfortunately, the expression data for ACE2 did not pass our quality filtering criteria even if it constitutes the main point of entry for viral infection of cells and would therefore, at least theoretically, configure an important candidate gene in these analyses. Spatial correlations between their expression profiles and structural brain deviations may help to explain region-specific vulnerability and support the hypothesis that SARS-CoV-2-related neuropathology is shaped by underlying molecular architecture.

### Cellular maps

To investigate the cellular substrates underlying the regional brain structural deviations observed in post-COVID syndrome, we derived cortical and subcortical cell-type distribution maps using the Bretigea package^47^. This analytical framework utilizes gene expression data from the Allen Human Brain Atlas (AHBA) and integrates curated gene sets derived from human single-cell transcriptomic studies to estimate the relative abundance of specific cell types across brain regions. The cell populations included in the analysis encompassed neurons, astrocytes, microglia, oligodendrocytes, oligodendrocyte precursor cells (OPCs), and endothelial cells. Gene markers corresponding to each cell class were used to compute regional enrichment scores, which were then aggregated and spatially aligned to the same Desikan-Killiany atlas used for our cortical thickness and surface area analyses. This ensured that cell-type density estimates were matched to the spatial resolution of our imaging data. We then assessed the relationship between cell-type densities and neuroanatomical deviations derived from the normative modelling framework. Specifically, we calculated region-wise correlations between cell-type density maps and z-scored deviations in cortical thickness and surface area. This approach would allow us to identify which cellular populations most strongly aligned with the pattern of structural brain changes observed in PCS, which in turn might hint at possible pathophysiological processes, including neuroinflammation (via microglial and astrocytic involvement), demyelination or oligodendrocyte dysfunction, and disruptions in neurovascular integrity.

### Neurotransmitter molecular targets

To further investigate the molecular architecture associated with regional brain alterations, we incorporated receptor density maps derived from normative Positron Emission Tomography (PET) imaging data. These templates represent in vivo estimates of neurotransmitter receptor and transporter availability across the human brain and were sourced from publicly available datasets generated using high-affinity radioligands in healthy adult populations and are available as part of the *neuromaps* toolbox^14^. The receptor maps included distributions for key molecular systems implicated in neuropsychiatric function and fatigue regulation, such as the serotonergic system (5-HT1A, 5-HT1B, 5-HT2A, 5-HT4, 5-HT6, 5HT transporter), dopaminergic system (D1, D2, and dopamine transporter [DAT]), glutamatergic system (NMDA, mGluR5), cholinergic system (M1 muscarinic receptor, vesicular acetylcholine transporter [VAChT]), cannabinoid system (CB1 receptor), GABAergic system (GABA-A receptor), opioid system (mu, kappa, and delta opioid receptors), noradrenergic system (norepinephrine transporter [NET]), and histaminergic system (H3 receptor). For details on the data originating each of these maps, we direct the reader to the original *neuromaps* publication^14^. Each PET-derived receptor density map was spatially registered and parcellated to match the Desikan-Killiany atlas used for cortical morphometry analyses. This alignment enabled region-wise comparison between individual structural deviations (e.g., cortical thickness or surface area z-scores from normative modelling) and normative receptor distributions. Associations were quantified using correlation and canonical correlation analyses, allowing us to assess which neurochemical systems best predicted the spatial distribution of observed structural alterations in PCS. This receptor-informed analysis would provide a neurochemical dimension to the observed anatomical deviations, facilitating the prioritisation of specific molecular systems that may underlie or modulate the persistent fatigue and neuropsychiatric symptoms reported in post-COVID syndrome.

### Statistical analysis

Statistical analyses were performed to examine group differences and associations across demographic, clinical, cognitive, and neuroimaging measures. Between-group comparisons for continuous demographic and clinical variables were conducted using independent-samples t-tests, with effect sizes reported as Cohen’s d. Categorical variables were compared using chi-square tests. Bayes factors (BF[[) were also computed to quantify the strength of evidence supporting group differences. For neuroimaging data, all group comparisons - including regional structural deviations (z-scores for cortical thickness, surface area, and subcortical volumes) - were evaluated using analysis of covariance (ANCOVA) models. These models included age, gender, handedness (manual dexterity), and total intracranial volume (TIV) as covariates to control for known sources of anatomical and physiological variability. To evaluate relationships between regional neuroanatomical deviations and molecular or cellular variables (candidate gene expression, cellular densities, receptor distributions), Spearman’s rank correlation analyses were conducted. Given the inherent spatial autocorrelation in cortical neuroimaging data, we employed permutation-based spin tests (1,000 permutations), generating null distributions by systematically rotating cortical maps while preserving spatial relationships (Vasa method)^48^. This rigorous approach ensured accurate assessments of statistical significance and minimized false-positive findings due to spatial dependencies. For multivariate integration of structural imaging metrics with molecular and cellular profiles, canonical correlation analysis (CCA) was performed ^49^. Statistical significance of canonical correlations was assessed through permutation testing (1,000 permutations), also accounting for spatial autocorrelation using spin permutations. Network-diffusion modelling outcomes were validated against null models generated by permutations of structural connectivity matrices (scrambled connectomes) and Euclidean-distance-based surrogate connectomes (1,000 permutations), further ensuring the robustness and specificity of observed diffusion patterns^50^. All statistical procedures and significance testing were performed using Python and R software packages specialized for neuroimaging and spatial analyses, with an alpha threshold set at p < 0.05 (two-tailed) for statistical significance, unless otherwise specified.

## Results

### Sociodemographics and clinical characterisation

Results have been first reported elsewhere. Here we describe the main findings that might be relevant to interpret the new neuroimaging findings. The groups were matched for age, sex, ethnicity, and body mass index (BMI), with no significant differences observed across these variables (all p > 0.5; Table 1). However, groups differed significantly in years of education, with HC participants reporting higher educational attainment. PCS participants also had a significantly longer duration since first SARS-CoV-2 infection, and were significantly less likely to have been vaccinated prior to infection. Clinically, individuals with PCS reported significantly greater symptom burden across multiple domains. These included fatigue (both visual analogue scale and multidimensional components), sleep disturbance (PSQI), post-exertional malaise, autonomic dysfunction (COMPASS-31), affective symptoms (HADS depression/anxiety), PTSD-related symptoms, and musculoskeletal complaints (WPI and SS scores). Notably, 40% of PCS participants met criteria for fibromyalgia, compared to none in the HC group. These differences were supported by both frequentist (p < 0.05) and Bayesian inference, with many outcomes yielding strong to extreme evidence in favor of group separation (BF□□ > 10), particularly for fatigue, post-exertional malaise, and pain sensitivity (Supplementary Table S1).

**Table 1.** Sociodemographic and clinical characteristics of PCS and healthy control participants. Group comparisons of demographic and COVID-19-related variables between individuals with Post-COVID-19 Syndrome (PCS) and matched healthy controls (HC). Data are shown as mean (standard deviation) or counts. Between-group differences were assessed using independent samples t-tests or chi-squared tests as appropriate. Bayes factors (BF□□) quantify evidence for group differences; values >1 indicate increasing support for the alternative hypothesis. Asterisks (*) denote statistically significant results at p < 0.05 or BF□□ > 1.

| Variable | PCS Mean (std) | HC Mean (std) | Statistics | p | B<sub>10</sub> |
| --- | --- | --- | --- | --- | --- |
| Age | 40.6 (10.9) | 40.0 (10.4) | T(38)=0.208 | 0.837 | 0.312 |
| Gender | 18F/2M | 17F/3M | $\chi^2(1)=0.229$ | 0.663 | N/A |
| Ethnicity | White: 18<br>Mixed: 0<br>Asian: 1<br>Black: 0<br>South Asian: 1<br>Hispanic: 0 | White: 15<br>Mixed: 1<br>Asian: 1<br>Black: 2<br>South Asian: 0<br>Hispanic: 1 | $\chi^2(5)=5.273$ | 0.384 | N/A |
| BMI | 23.8 (3.44) | 24.4 (3.84) | T(38)=-0.55 | 0.582 | 0.346 |
| Years of education | 17.30 (2.94) | 19.35 (3.13) | T(38)=-2.13 | 0.039* | 1.845 |
| Time since first infection (months) | 23.0 (9.17) | 15.7 (11.5) | T(38)=2.21 | 0.033* | 0.749 |
| Vaccination prior infection | Yes: 4<br>No: 16 | Yes: 14<br>No: 6 | $\chi^2(1)=10.101$ | 0.001* | N/A |

### Clinical routine blood tests

Group comparisons of routine blood parameters revealed no major differences between PCS and HC participants (Supplementary Table S2). However, PCS participants showed numerically higher creatinine and Gamma-GT levels, with corresponding Bayes factors (BF□□ = 1.232 and 1.735, respectively) providing anecdotal to moderate evidence for group differences. Platelet count also trended higher in the PCS group (p = 0.08; BF□□ = 1.123). Other metabolic, inflammatory, and haematological markers - including C-reactive protein, white and red cell counts, haemoglobin, and thyroid function - did not differ significantly between groups.

### Serum cytokines and glial markers

No statistically significant differences were observed in circulating levels of IFN-γ, IL-1β, IL-6, IL-8, IL-10, IL-13, TNF-α or GFAP (Supplementary Table S3). However, TNF-α (p = 0.116; BF□□ = 1.094) and S100β (p = 0.149; BF□□ = 1.025) showed suggestive evidence of elevation in the PCS group, consistent with mild glial or immune activation.

### Cognitive performance

Cognitive testing using the Cognitron battery revealed subtle but functionally relevant differences between groups (Supplementary Table S4). PCS participants showed significantly poorer performance on the delayed object memory task (RT_DFE; p = 0.02; BF□□ = 2.653), as well as trend-level impairments in the Lead Balloon task (abstract reasoning; p = 0.06; BF□□ = 1.330) and vigilance (Spotter task RT; p = 0.06; BF□□ = 1.303). Other domains - including motor control, 2D spatial manipulation, and verbal analogies - did not show significant differences. Overall, Bayesian analysis provided moderate evidence for reduced memory and attentional function in PCS.

### Group differences in the total amounts of positive and negative extreme deviations

Using our predefined threshold of |z|>1.96, extreme deviations were mostly present for thickness and absent for surface and subcortical volume. PCS and control participants did not differ in the total number of either positive or negative extreme deviations in cortical thickness (positive: F(35)=0.082, p=0.776; negative: F(35)=0.368, p=0.548).

**Figure 1.**
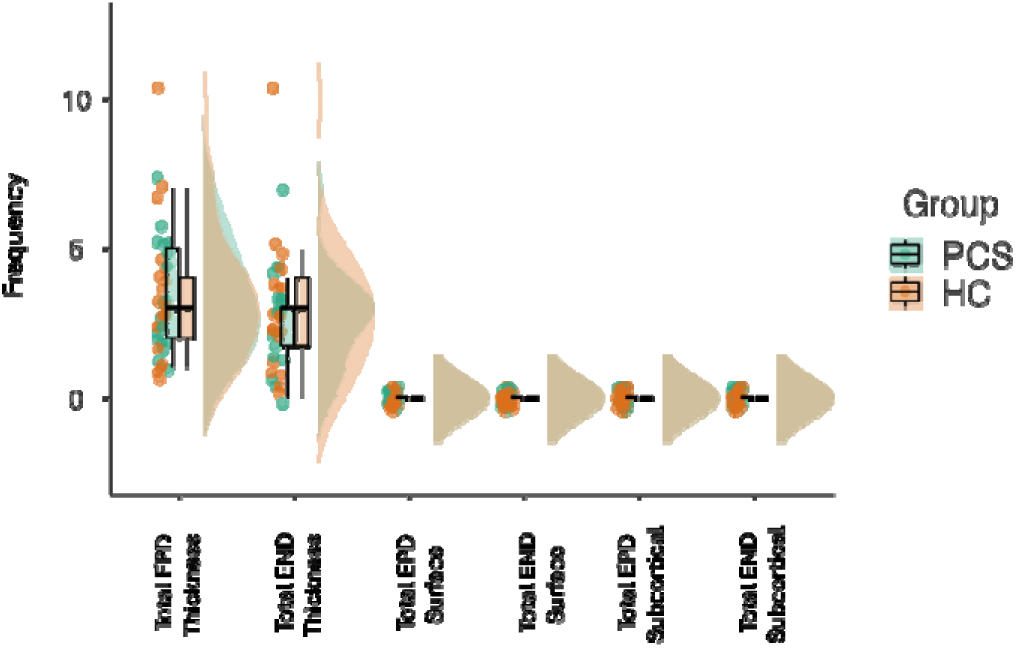
Distribution of Individual-Level Extreme Neuroanatomical Deviations in PCS and Healthy Controls. Violin plots showing the distribution of total extreme deviations in cortical and subcortical morphometry metrics by group (PCS vs HC). Each plot displays the number of extreme deviations per participant (z-scores |z| > 1.96) across six categories: positive and negative deviations in cortical thickness, surface area, and subcortical volume. Violin widths reflect kernel density estimates, with overlaid boxplots showing medians and interquartile ranges. Individual data points are also plotted. PCS = post-COVID syndrome group; HC = healthy controls; EPD – extreme positive deviations; END – extreme negative deviations.

### Links between total amounts of positive and negative extreme thickness deviations and clinical, blood and cognitive parameters

Exploratory partial spearman correlations accounting for group, age, gender, dexterity and TIV revealed significant positive correlations between the total amount of positive extreme thickness deviations and performance in the Spotter task and blood magnesium, and negative correlations with AST levels, mental fatigue and total FAI scores, as well as severity of autonomic and immune symptoms assessed by the DePaul questionnaire. In the case of the total amount of negative extreme thickness deviations we found positive associations with serum S100β, blood creatinine and urea, and blood count of eosinophils. None of these associations would have survived FDR correction for the total number of parameters assessed (Supplementary Table S5).

### Group differences in regional structural deviation Z-scores

Normative modelling revealed subtle but spatially distinct patterns of cortical deviation in PCS. At the group level, patients exhibited reduced cortical thickness deviation Z-scores in the left medial and right lateral orbitofrontal cortex, right pars opercularis and triangularis of the frontal cortex and left fusiform gyrus; and increased deviation Z-scores thickness in the left lateroccipital cortex and right post-central gyrus (Figure 2). Surface area Z-score deviation scores showed reduced values in the right postcentral gyrus and increases in the right frontal pole. None of these regions would have survived FDR correction for the number of regions tested. For subcortical volumes, no comparisons reached significance.

**Figure 2.**
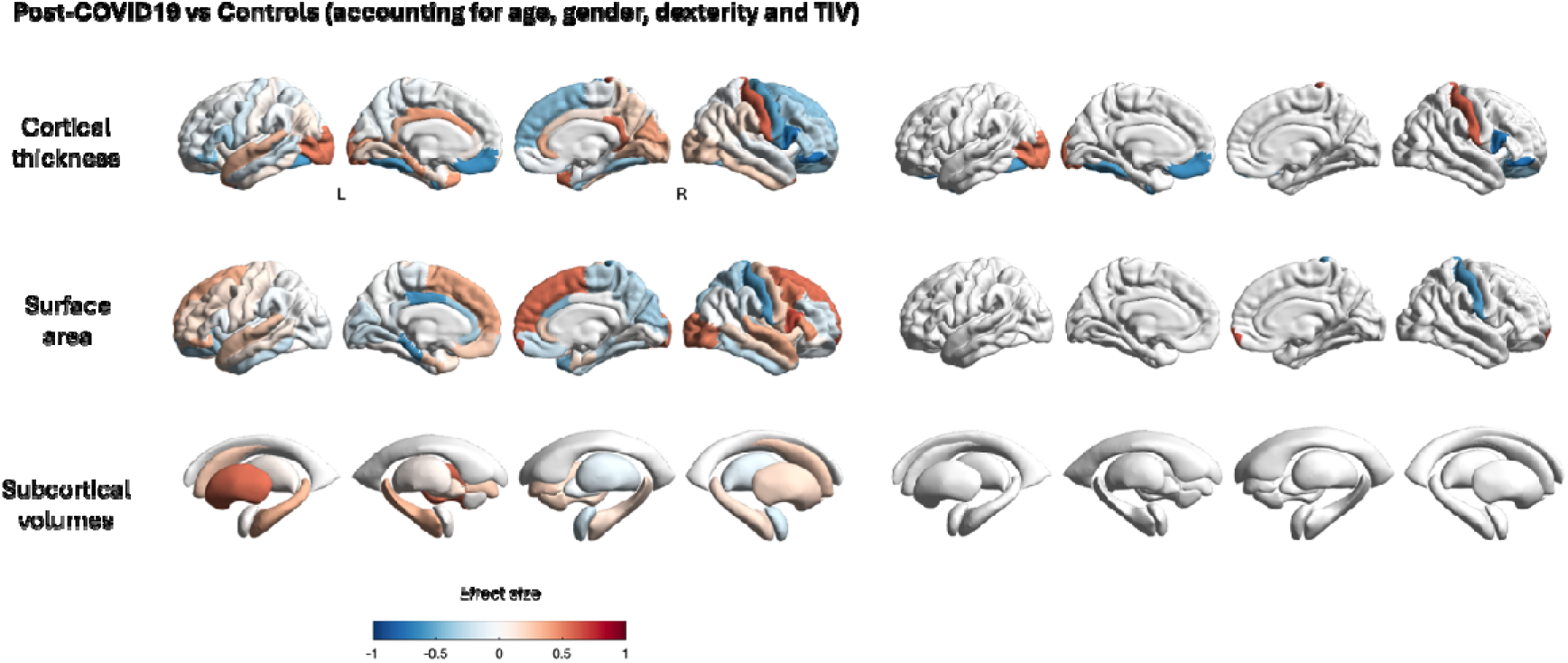
Regional differences in cortical thickness, surface area, and subcortical volumes deviation scores between post-COVID-19 patients and healthy controls. Comparisons were adjusted for age, gender, handedness, and total intracranial volume (TIV). Colours indicate effect sizes (Cohen’s d), with red representing increased measures and blue indicating decreased measures in post-COVID-19 patients compared to controls. Significant results are thresholded at uncorrected p < 0.05.

As a sensitivity analysis and because alterations in brain structure in PCS are often simply attributed to the presence of olfactory dysfunction, we repeated the same group comparisons on deviation Z-scores this time additionally accounting for the presence of anosmia/ageusia. As shown in Supplementary Figure S1, many of the reported group differences were still evident and others not reaching significance before did after this additional covariate. For cortical thickness, we found decreases in Z scores for left fusiform and middle orbitofrontal gyri and an increase for the right isthmus cingulate. For surface area, we found increases in the left caudal middle frontal gyrus, left pars orbitalis and right pars triangularis of the frontal gyrus, and right frontal pole, and decreases in the right postcentral gyrus (Supplementary Table S6).

### Regional and circuit overlap of extreme positive and negative deviations

Extreme deviations were heterogeneous at the individual regional level with every individual region being identified as extreme in <35% of the members of either group. However, a circuit-based overlap analysis, where we considered deviated regions as well as all other brain regions structurally connected to them, revealed significant convergence within specific structural networks, affecting up to an excess of 50% of patients in the PCS group. These deviations mapped predominantly onto salience and attention-related circuits, suggesting distributed but coherent alterations in anatomically and functionally connected regions (Figure 3).

**Figure 3.**
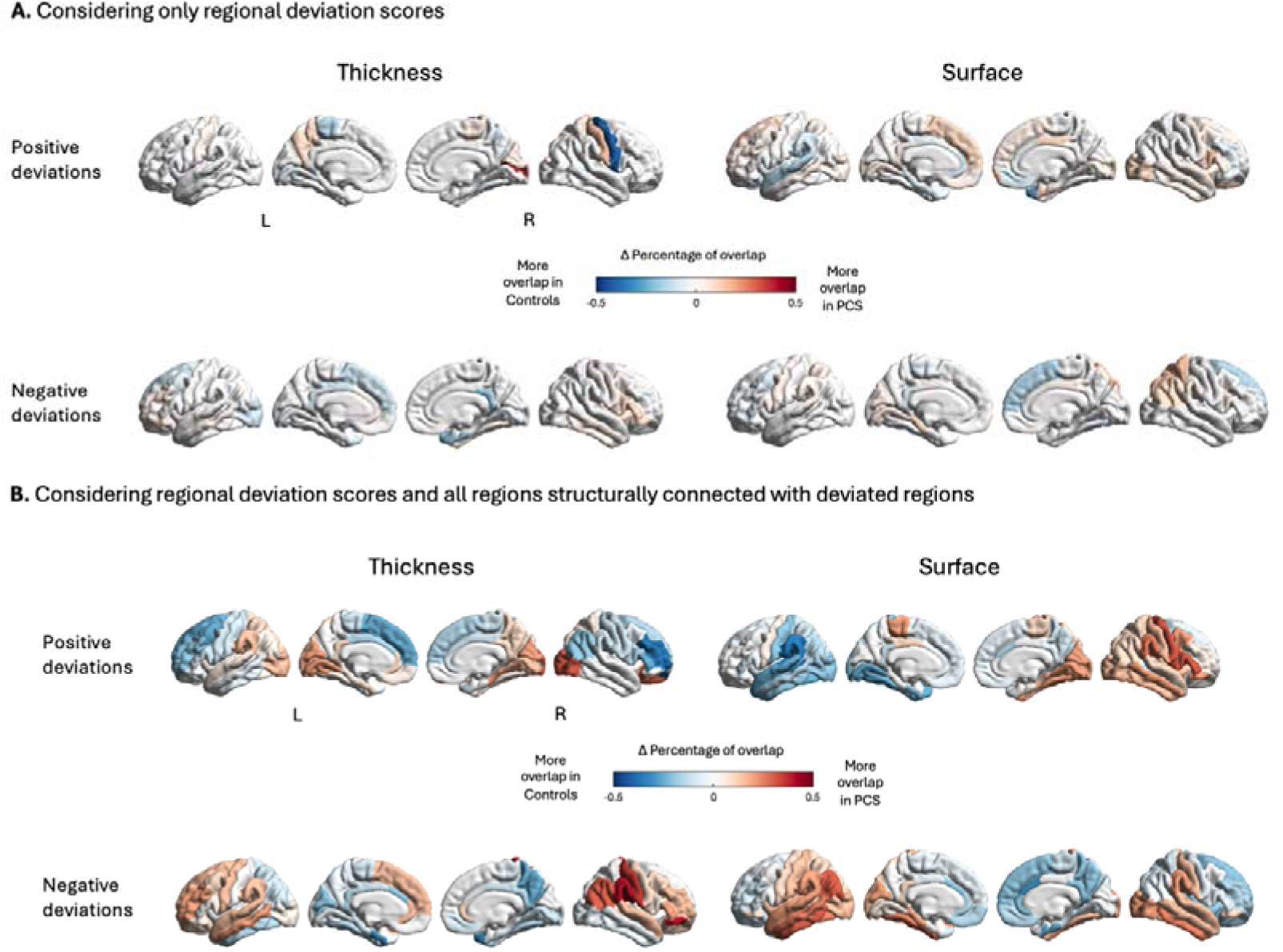
Regional (A) and circuit-based (B) overlap analysis comparing positive and negative deviations in cortical thickness and surface area between post-COVID syndrome (PCS) patients and healthy controls. Colour gradients indicate the difference (Δ percentage overlap) in the proportion of participants showing deviations, with red reflecting greater overlap in PCS patients and blue representing greater overlap in controls.

### Network-Diffusion Modelling

Network diffusion modelling demonstrated that structural deviations in PCS are not randomly distributed but follow pathways consistent with structural connectivity. The optimized model (β = 0.010) produced simulated deviation patterns that significantly correlated with empirical data (Figure 4). Posterior-temporoparietal regions emerged as likely “seeds” for network-wide propagation, showing the highest seed likelihood scores. These regions are involved in multisensory integration and attention and may represent key vulnerability nodes in PCS. Permutation testing confirmed that these findings exceeded chance expectations (p < 0.05), with simulations based on scrambled connectomes and Euclidean distance metrics showing significantly lower correspondence with empirical data.

**Figure 4.**
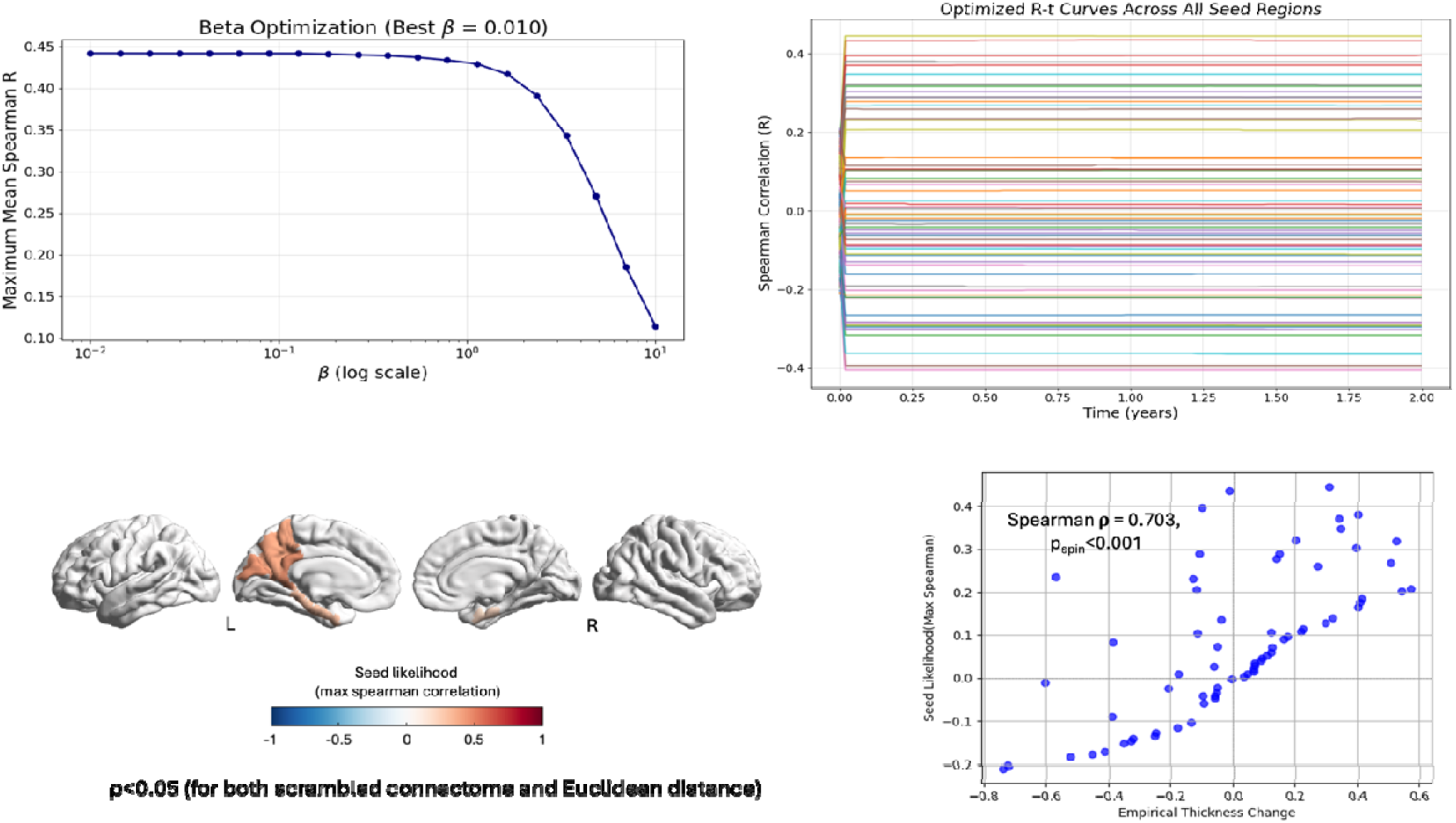
Network diffusion modelling results. Upper left panel shows beta optimization, identifying the best diffusion coefficient (β = 0.010) based on maximum mean Spearman correlation. Upper right panel illustrates optimized Spearman correlation (R) curves over time across all potential seed regions. The lower left panel maps seed likelihood (maximum Spearman correlation) across brain regions, with higher likelihood regions indicated in red. The lower right panel demonstrates the significant positive correlation between seed likelihood and empirical cortical thickness changes (Spearman ρ = 0.703, p < 0.001). Permutation testing against scrambled connectomes and Euclidean distance models confirmed the significance of these findings (p < 0.05).

### Candidate genes related to SARS-CoV-2 entry into host cells

Spatial correlations between neuroanatomical deviations and candidate gene expression revealed strong associations with TMPRSS2, a known facilitator of SARS-CoV-2 entry into host cells. TMPRSS2 expression was significantly associated with regional changes in cortical thickness, surface area, and seed likelihood from network diffusion models (Figure 5). Other entry-related genes, such as FURIN, NRP1, and TLR4, showed weaker or nonsignificant associations.

**Figure 5.**
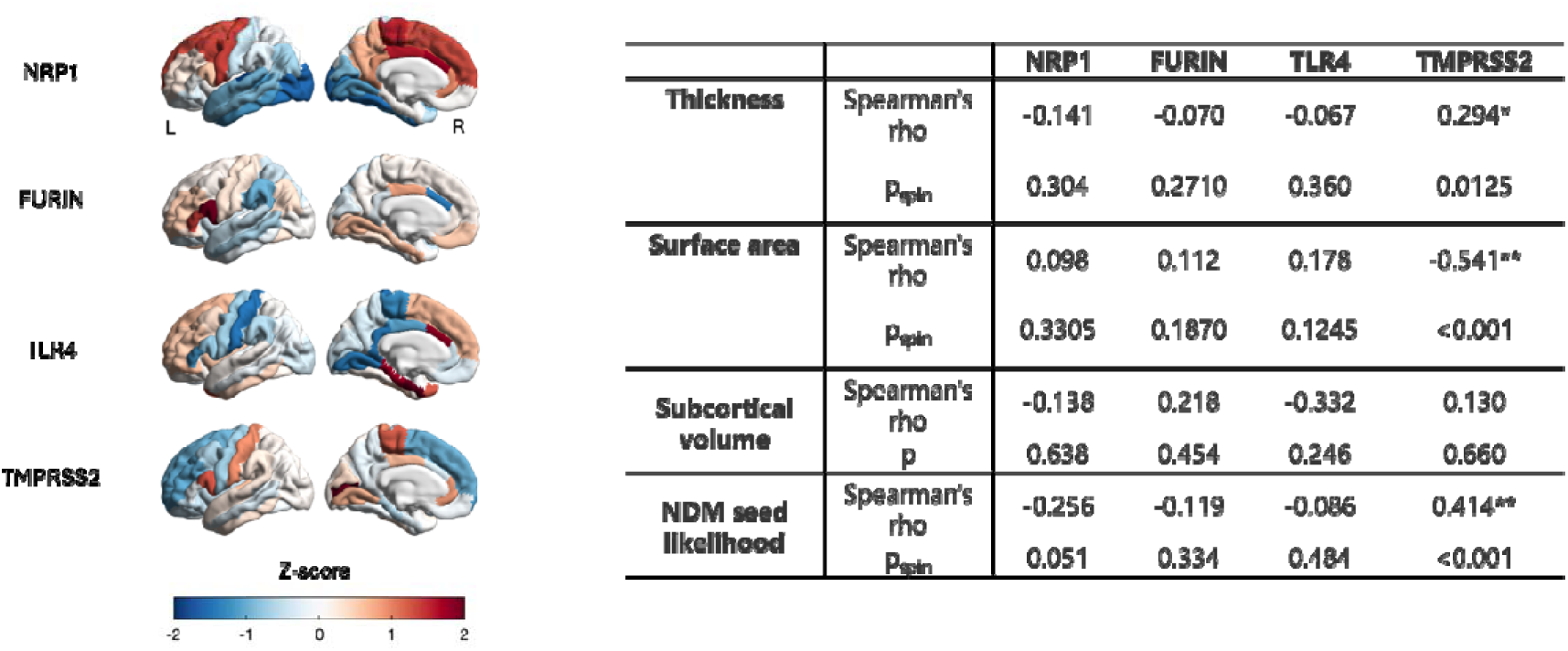
Associations between SARS-CoV-2-related gene expression (NRP1, FURIN, TLR4, TMPRSS2) and neuroimaging metrics (cortical thickness, surface area, subcortical volume, and network diffusion model [NDM] seed likelihood). Left panel illustrates regional mRNA expression (Z-scores). Right panel summarizes Spearman’s correlations (rho) and associated permutation-based spin-test p-values (p_spin_), highlighting significant associations (*p<0.05, \*\**p<0.001)*.

### Cellular and Neurochemical Correlates

Canonical correlation analyses (CCA) identified significant relationships between brain structural deviations and regional cell-type distributions. The strongest loadings involved neurons and microglia, suggesting a role for neuroinflammatory and excitatory/inhibitory imbalances in PCS-related alterations (Figure 6A). Associations were stronger for cortical thickness than surface area, consistent with prior work showing thickness as a sensitive index of neuroimmune changes.

Receptor-based CCA further linked cortical deviations in thickness with a multivariate pattern of neurotransmitter systems. The most robust negative associations were observed for serotonergic 5HT1A and B, 5HT6, cholinergic (VAChT), and cannabinoid (CB1) targets, while mGlu5R had the highest positive loading (Figure 6B). The neurotransmitter system CCA model did not reach significance for deviation in surface area.

**Figure 6.**
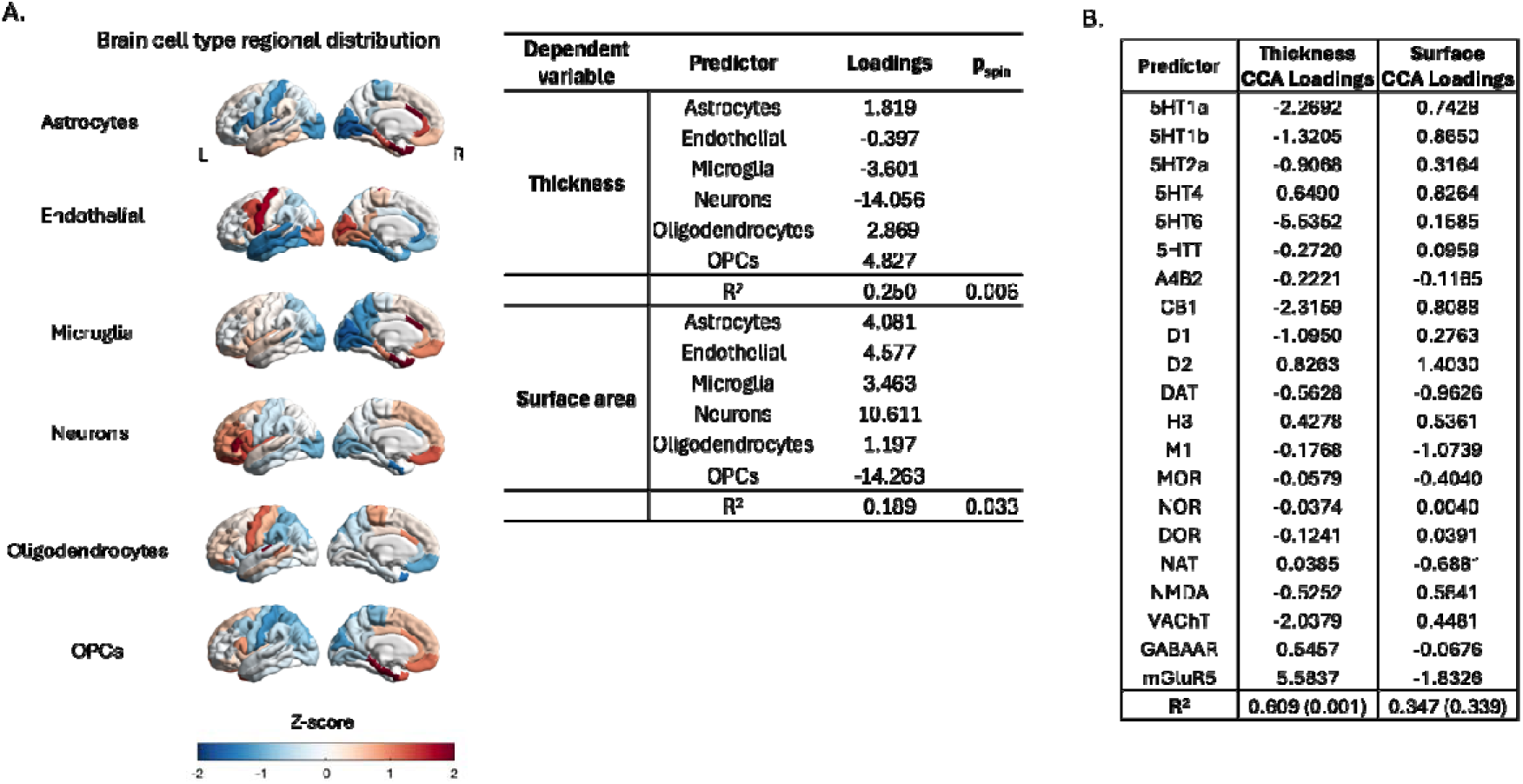
Cellular and neuroreceptor correlates of structural deviations in PCS. (A) Brain cell-type regional distributions illustrating Z-scores for astrocytes, endothelial cells, microglia, neurons, oligodendrocytes, and oligodendrocyte precursor cells (OPCs). The accompanying table summarizes canonical correlation analysis (CCA) loadings and permutation-based spin-test p-values (p_spin_), indicating significant associations between specific cell types and regional alterations in cortical thickness and surface area. (B) Canonical correlation analysis loadings for neuroreceptor densities predicting cortical thickness and surface area alterations. Predictors include serotonergic (5HT receptors), cannabinoid (CB1), dopaminergic (D1, D2, DAT), histaminergic (H3), cholinergic (M1, VAChT), glutamatergic (NMDA, mGluR5), opioid (MOR, NOR, DOR), noradrenergic (NAT), amyloid-beta (A4B2), and GABAergic (GABAAR) systems. The table also reports R² values with corresponding permutation-based significance (p-values in parentheses), highlighting stronger associations for thickness compared to surface area.

## Discussion

We applied a multimodal analytic framework to investigate neuroanatomical deviations in individuals with PCS and persistent fatigue following mild SARS-CoV-2 infection. By integrating individualized morphometric deviation mapping with gene expression profiles, cellular composition, receptor distributions, and network diffusion modelling, we identified spatially organized patterns of cortical change. Although individual regional deviations were modest and variable, they converged across structurally connected circuits and aligned with specific molecular and neurochemical features. While preliminary, these findings lay the groundwork for future studies seeking to clarify the biological basis of post-COVID fatigue and identify potential targets for intervention.

We observed reduced cortical thickness in orbitofrontal regions and increased thickness in occipital and sensory cortices. These changes align with prior evidence of frontal vulnerability in chronic fatigue and post-infectious conditions and may reflect disruption of circuits involved in reward processing, emotional regulation, and sensory integration^6–8^. Rather than affecting the same regions uniformly, deviations clustered within structurally connected networks, suggesting that circuit-level organization better accounts for symptom-related alterations than isolated anatomical loci. Similar conclusions have been reached regarding brain structure and function within the context of other classical neuropsychiatric disorders^40^.

Network diffusion modelling extended these findings by identifying posterior-temporoparietal regions as likely sources for a slow propagation of structural alterations across the connectome, reinforcing their role as integrative hubs for sensory and attentional information - functions frequently disrupted in PCS^1, 51^ ^52, 53^. We optimized a diffusion coefficient (β = 0.010) within a biologically realistic range, indicating a slow and spatially constrained propagation pattern^19–23^. The same regions also showed higher expression of TMPRSS2, a host gene required for SARS-CoV-2 cellular entry^44^. While TMPRSS2 expression alone does not imply ongoing viral effects, its spatial correlation with structural deviation patterns suggests that pre-existing molecular architecture may influence regional vulnerability during or after SARS-CoV-2 infection^54, 55^.

We found that cortical thickness deviations correlated with normative maps of neuronal and microglial density. Regions with greater neuronal density may be more susceptible to deviation through increased metabolic or synaptic demands^56, 57^. Microglial alignment may reflect structural consequences of immune surveillance or low-grade inflammation^58, 59^. These associations highlight potential cellular substrates of PCS-related cortical changes. In parallel, canonical correlation analyses linked structural deviations to the distribution of serotonergic, cholinergic, cannabinoid, and glutamatergic receptors. Regions enriched in 5-HT1A, 5-HT1B, 5-HT6, VAChT, CB1, and mGlu5 showed the strongest anatomical correspondence. These systems contribute to arousal, mood, cognition, and fatigue regulation^60–64^. While we did not assess receptor binding or neurotransmitter function directly, the spatial alignment with structural deviations indicates that cortical regions embedded in specific neuromodulatory environments might be differentially affected in PCS. These findings are particularly interesting within the context of ongoing pharmacological trials investigating neuromodulatory agents - such as vortioxetine, which modulates 5-HT1A, 5-HT1B, and 5-HT6 receptors^65^ - for the treatment of cognitive and affective symptoms in PCS^66^.

PCS participants showed impairments in delayed visual memory and sustained attention, with slower reaction times on delayed object recall and trend-level reductions in vigilance performance. These cognitive deficits were domain-specific and aligned with structural deviations in circuits supporting attention and episodic memory. Notably, sustained attention performance correlated with the burden of extreme positive deviations in cortical thickness, suggesting that individuals showing greater anatomical deviation also exhibited measurable functional change^35, 67^. Although serum biomarkers showed no major group-level differences, exploratory analyses revealed associations between cortical thinning and S100β, creatinine, urea, and eosinophils, consistent with subtle glial and systemic contributions to brain structure. These patterns converge on a model in which regional structural alterations, likely embedded within neuroimmune and neuromodulatory gradients, underlie selective cognitive dysfunction in PCS.

Our study’s integrative approach - combining personalized structural modelling with gene expression and receptor data - represents a methodological advance and supports a biologically coherent narrative of PCS-related fatigue. Nevertheless, several limitations must be acknowledged. This study has important limitations. The sample size was modest, limiting power and precision of multivariate associations. The cross-sectional design precludes inference about temporal progression or reversibility. While we adjusted for key demographic and imaging covariates, unmeasured confounders - such as variations in SARS-COV2 variant and off-label treatment with supplements/hormonal therapy/histamine blockers - may have influenced both brain structure and symptom expression. Molecular and receptor associations were based on population-average postmortem datasets - including the Allen Human Brain Atlas, which includes only six adult donors - limiting generalizability and precluding inferences about individual-level or state-dependent biology^68^. The receptor maps derived from PET templates also reflect healthy population averages and do not capture potential dynamic changes in neurotransmitter function following infection/inflammation^69^.

Taken together, our findings reveal that PCS-related fatigue is associated with distributed cortical deviations that follow structural connectivity and co-localize with specific molecular and cellular features. These results characterize PCS as a condition involving spatially structured brain alterations embedded within established neurochemical and immune architectures. Future research should aim to replicate and extend these findings in larger, longitudinal cohorts, ideally integrating multimodal imaging with direct measures of neuroinflammation and neurotransmitter function.

## Credit authorship statement

**Daniel Martins**: Conceptualization, Methodology, Formal analysis, Investigation, Writing – original draft, Supervision, Project administration. **Ziyuan Cai**: Data processing, Quality control, Writing – review & editing. **Nicole Mariani**: Methodological support, Writing – review & editing. **Alessandra Borsini**: Methodological support, Writing – review & editing. **Valeria Mondelli**: Conceptualization, Writing – review & editing. **Brandi Eiff**: Participant recruitment, Investigation, Data curation. **Silvia Rota**: Participant recruitment, Clinical assessments, Project coordination, Writing – review & editing. **Daniel van Wamelen**: Participant recruitment, Writing – review & editing. **Timothy Nicholson**: Clinical oversight, Participant recruitment, Writing – review & editing. **Aleksandra Podlewska**: Setup, Project coordination, Writing – review & editing. **Ray Chaudhuri:** Setup, Project coordination, Writing – review & editing. **Laila Raida**: Cognitive tasks setup, Writing – review & editing. **Adam Hampshire**: Cognitive task development, Normative modelling, Software, Writing – review & editing. **Federico Turkheimer**: Statistical supervision, Interpretation, Writing – review & editing. **Steven C. R. Williams**: Resources, Funding acquisition, Supervision, Writing – review & editing. **Fernando Zelaya**: Conceptualization, Methodology, Supervision, Writing – review & editing. **Mattia Veronese**: Methodology, Supervision, Writing – review & editing. All authors have read and approved the final manuscript and agree to be accountable for all aspects of the work.

## Acknowledgements

This study was funded by the National Institute for Health and Care Research (NIHR) Maudsley Biomedical Research Centre (BRC), South London and Maudsley NHS Foundation Trust, under the “Reach Out” funding call (Ref: R0-01). We are grateful to all participants for their time and commitment to the study. We thank the radiographers and technical staff at the Centre for Neuroimaging Sciences (King’s College London) for their assistance with MRI acquisition. We also acknowledge Synnovis (Viapath) for processing routine blood analyses, the Clinical Research Facility (CRF) team at King’s College Hospital Foundation Trust for their support during participant visits and blood collection, and the Cognitron platform team for assistance with cognitive data management and normative modeling.

