## supplementary for "Molecular, cellular and network mapping of brain structural deviations in patients with Post-COVID19 syndrome"

**Supplementary Table S1. Self-reported fatigue, affective symptoms, and somatic complaints in PCS and HC participants, with Bayesian inference.** Comparison of symptom severity between individuals with Post-COVID-19 Syndrome (PCS) and matched healthy controls (HC) across domains including fatigue (VAS, MFI), sleep (PSQI), autonomic function (COMPASS-31), post-exertional malaise, mood (HADS), PTSD, and fibromyalgia-related symptoms. Values are shown as mean (standard deviation), and group comparisons were conducted using independent samples t-tests or chi-squared tests, as appropriate. The Bayesian factor (BF₁₀) is reported for each comparison, quantifying the strength of evidence in favor of the alternative hypothesis. Asterisks (*) denote statistically significant differences at p < 0.05 or BF₁₀ > 1.

| **Variable** | **PCS**  **Mean (std)** | **HC**  **Mean (std)** | **Statistics** | **P-value** | **B_10_** |
| --- | --- | --- | --- | --- | --- |
| **Fatigue VAS** | 5.500 (1.606) | 1.850 (1.663) | T(38)=7.061 | 2.036x10^-8^* | 876.874* |
| **Multidimensional fatigue**  **General** **Physical**  **Reduced activity**  **Reduced motivation**  **Mental fatigue** | 15.200 (3.205)  14.600 (2.998)  12.150 (3.703)  8.600 (3.393)  12.850 (3.843) | 6.684 (2.212)  5.737 (2.884)  5.444 (2.307)  4.895 (2.378)  6.053 (3.027) | T(38)=47.68  T(38)=43.19  T(38)=15.09  T(38)=9.59  T(38)=21.41 | 1.355x10^-11*^  2.408x10^-11^*  1.072x10^-7*^  3.589x10^-4^*  4.391x10^-7^* | 1332.772*  4158.129*  650.746*  34.338*  767.860* |
| **ESS** | 12.400 (5.548) | 6.474 (4.234) | T(37)=3.735 | 6.304x10^-4^* | 24.874* |
| **HADS Depression** | 8.900 (2.954) | 4.053 (2.345) | T(37)=5.657 | 1.833x10^-6^* | 153.950* |
| **HADS Anxiety** | 7.550 (3.790) | 5.368 (2.477) | T(37)=2.116 | 0.041* | 1.287* |
| **PTSD total score** | 19.750 (10.208) | 6.105 (8.185) | T(37)=4.590 | 4.964x10^-5*^ | 59.715* |
| **De Paul's**  **PEM** | 2.950 (0.945) | 0.421 (0.449) | T(37)=10.584 | 9.541x10^-13*^ | 64546.865* |
| **De Paul's**  **Unrefreshing sleep** | 3.400 (0.821) | 1.316 (0.749) | T(37)=8.269 | 6.229x10^-10*^ | 1547.301* |
| **De Paul's**  **Autonomic symptoms** | 1.775 (0.881) | 0.553 (0.550) | T(37)=5.166 | 8.438x10^-6*^ | 73.625* |
| **De Paul's**  **Neurocognitive** | 1.775 (0.910) | 0.474 (0.612) | T(37)=5.212 | 7.316x10^-6*^ | 401.197* |
| **De Paul's**  **Immune** | 2.250 (0.993) | 0.658 (0.834) | T(37)=5.405 | 4.010x10^-6*^ | 381.905* |
| **COMPASS total score** | 62.413 (19.629) | 30.666 (19.158) | T(37)=5.108 | 1.011x10^-5*^ | 569.590* |
| **Global PSQI** | 15.200 (6.542) | 9.263 (5.791) | T(37)=2.995 | 0.005* | 15.254* |
| **MRC**  **Dyspnea**  **Scale** | Stage 1: 9  Stage 2: 9  Stage 3: 2 | 19  1  0 | χ^2^(5)=11.971 | 0.003* | N/A |
| **MoCA** | 26.8 (2.55) | 26.5 (1.54) | T(37)=0.430 | 0.668 | 0.420 |
| **Pain detect**  **WPI**  **SS** | 9.00(6.53)  5.45 (4.07)  8.55 (1.88) | 1.7 (1.84)  1.00 (1.62)  1.75 (1.68) | T(37)= 4.813  T(37)=4.541  T(37)=12.066 | <0.001*  <0.001*  <0.001* | 1315.956*  119.442*  13663.606* |
| **Fibromyalgia**  **diagnostic criteria** | Yes: 8  No: 12 | Yes:0  No:20 | χ^2^(2)=10.581 | 0.005* | N/A |

**Supplementary Table S2. Comparison of standard blood parameters between PCS and healthy control participants.** Descriptive statistics and group comparisons for hematological, biochemical, and inflammatory markers in individuals with Post-COVID-19 Syndrome (PCS) and matched healthy controls (HC). Results are reported as mean (standard deviation). Between-group comparisons were conducted using the Mann–Whitney U test (W), with corresponding p-values and Bayesian factors (BF₁₀) reported for each variable. Asterisks (*) denote statistically significant differences at p < 0.05 or BF₁₀ > 1.

| **Variable** | **PCS Mean (SD)** | **HC Mean (SD)** | **Statistics**  **W** | **P-value** | **B_10_** |
| --- | --- | --- | --- | --- | --- |
| **Sodium** | 140.118 (2.619) | 140.474 (2.091) | 161.00 | 1.00 | 0.369 |
| **Potassium** | 3.965 (0.237) | 4.000 (0.252) | 126.00 | 0.53 | 0.334 |
| **Urea** | 4.135 (1.346) | 4.337 (0.942) | 158.50 | 0.94 | 0.359 |
| **Creatinine** | 61.235 (8.318) | 68.632 (11.847) | 101.50 | 0.06 | 1.232* |
| **Glucose** | 4.871 (0.819) | 4.968 (1.106) | 157.00 | 0.90 | 0.329 |
| **Calcium_corrected** | 2.268 (0.059) | 2.288 (0.109) | 159.00 | 0.95 | 0.341 |
| **Phosphatase** | 1.088 (0.166) | 1.171 (0.163) | 106.00 | 0.08 | 0.749 |
| **Proteins_total** | 71.000 (3.691) | 71.526 (3.878) | 148.50 | 0.69 | 0.337 |
| **Albumin** | 47.000 (1.969) | 47.000 (2.749) | 154.50 | 0.83 | 0.326 |
| **Bilirubin_total** | 10.125 (6.672) | 9.824 (7.170) | 116.00 | 0.48 | 0.370 |
| **ALP** | 64.176 (15.204) | 63.947 (23.710) | 145.50 | 0.62 | 0.362 |
| **AST** | 23.353 (6.441) | 22.316 (7.521) | 131.00 | 0.34 | 0.420 |
| **Gamma_GT** | 13.059 (3.508) | 31.632 (47.978) | 97.50 | 0.04 | 1.735* |
| **Magnesium** | 0.859 (0.043) | 0.872 (0.054) | 151.50 | 0.76 | 0.347 |
| **CRP_recoded** | 1.353 (0.606) | 1.368 (0.597) | 158.50 | 0.92 | 0.397 |
| **Globulin** | 23.529 (3.875) | 24.526 (2.674) | 138.50 | 0.47 | 0.497 |
| **Ferritin** | 108.353 (103.511) | 99.789 (84.020) | 156.50 | 0.89 | 0.307 |
| **WCC** | 6.629 (1.662) | 6.501 (1.803) | 156.50 | 0.89 | 0.336 |
| **RCC** | 4.385 (0.457) | 4.561 (0.386) | 112.50 | 0.12 | 0.843 |
| **Hemoglobin** | 134.000 (13.224) | 137.263 (10.514) | 138.50 | 0.48 | 0.426 |
| **Haematocrit** | 0.403 (0.036) | 0.417 (0.029) | 120.00 | 0.19 | 0.739 |
| **MCV** | 91.988 (4.276) | 91.679 (5.684) | 159.00 | 0.95 | 0.329 |
| **MCH** | 30.624 (1.719) | 30.200 (2.149) | 151.00 | 0.75 | 0.348 |
| **MCHC** | 332.882 (11.931) | 329.158 (8.662) | 128.50 | 0.30 | 0.504 |
| **RDW** | 12.406 (0.599) | 12.474 (0.726) | 154.50 | 0.84 | 0.329 |
| **Platelet_count** | 267.118 (42.355) | 289.789 (45.281) | 105.00 | 0.08 | 1.123* |
| **MPV** | 10.988 (0.834) | 10.716 (0.727) | 121.00 | 0.20 | 0.463 |
| **Neutrophils** | 4.142 (1.178) | 3.773 (1.409) | 135.50 | 0.42 | 0.414 |
| **Lymphocytes** | 1.934 (0.550) | 2.029 (0.650) | 144.50 | 0.60 | 0.334 |
| **Monocytes** | 0.441 (0.102) | 0.441 (0.127) | 155.00 | 0.85 | 0.319 |
| **Eosinophils** | 0.141 (0.100) | 0.214 (0.243) | 147.50 | 0.67 | 0.365 |
| **Basophils** | 0.052 (0.030) | 0.042 (0.018) | 136.00 | 0.42 | 0.486 |
| **Immature_Granulocyte_Count** | 0.022 (0.016) | 0.022 (0.012) | 141.50 | 0.51 | 0.327 |
| **ESR** | 7.588 (6.662) | 8.278 (8.072) | 139.50 | 0.67 | 0.354 |
| **TSH** | 1.819 (0.630) | 1.633 (0.658) | 124.50 | 0.25 | 0.423 |
| **T4_free** | 15.124 (1.759) | 15.042 (1.624) | 156.00 | 0.87 | 0.321 |
| **NLR** | 2.209 (0.594) | 2.038 (1.137) | 125.50 | 0.26 | 0.540 |
| **MLR** | 0.234 (0.041) | 0.230 (0.072) | 156.00 | 0.87 | 0.346 |

**Supplementary Table S3. Serum cytokines and glial biomarkers in PCS and healthy control participants.** Group comparisons for circulating immune markers (interleukins, interferon-γ, TNF-α) and glial markers (S100β, GFAP) measured in serum samples from individuals with Post-COVID-19 Syndrome (PCS) and healthy controls (HC). Data are reported as mean (standard deviation). Concentrations are in pg/mL. Between-group differences were evaluated using the Mann–Whitney U test (W), with corresponding p-values and Bayes Factors (BF₁₀). Asterisks (*) indicate statistically suggestive effects (BF₁₀ > 1)

| **Variable** | **PCS Mean (SD)** | **HC Mean (SD)** | **Statistics**  **W** | **P-value** | **B_10_** |
| --- | --- | --- | --- | --- | --- |
| **IFNy** | 4.639 (8.032) | 4.182 (8.177) | 151.000 | 0.789 | 0.345 |
| **IL1b** | 0.345 (0.745) | 0.094 (0.061) | 195.000 | 0.300 | 0.573 |
| **IL6** | 1.197 (2.795) | 0.418 (0.205) | 194.000 | 0.478 | 0.431 |
| **TNFa** | 1.164 (1.799) | 0.980 (0.316) | 110.000 | 0.116 | 1.094* |
| **IL10** | 0.576 (1.145) | 0.229 (0.141) | 171.000 | 0.449 | 0.365 |
| **IL13** | 0.998 (0.889) | 1.228 (0.894) | 127.000 | 0.563 | 0.398 |
| **IL8** | 6.106 (4.671) | 6.581 (2.791) | 126.000 | 0.289 | 0.460 |
| **s100b** | 3.908 (2.192) | 5.165 (2.558) | 130.000 | 0.149 | 1.025* |
| **GFAP** | 14.62 (4.79) | 13.78 (5.32) | 157.000 | 0.515 | 0.406 |

**Supplementary Table S4. Performance on computerized cognitive tasks in PCS and healthy control participants.** Comparison of standardized deviation-from-expected (DFE) scores derived from the Cognitron cognitive task battery, assessing domains such as spatial reasoning, memory, motor control, and sustained attention. Scores represent z-standardized residuals relative to normative models controlling for age, sex, handedness, and ethnicity. Between-group differences were tested using independent samples t-tests, with corresponding p-values and Bayes Factors (BF₁₀). Asterisks (*) indicate results meeting the threshold for statistical significance (p < 0.05) or annedoctal Bayesian evidence (BF₁₀ > 1).

| **Variable** | **PCS Mean (SD)** | **HC Mean (SD)** | **Statistics** | **P-value** | **B_10_** |
| --- | --- | --- | --- | --- | --- |
| **Blocks_RT_DFE** | 0.136 (0.613) | -0.043 (0.539) | T(38)=0.981 | 0.33 | 0.452 |
| **BlocksSummaryScore_DFE** | -0.706 (1.069) | -0.301 (1.011) | T(38)=-1.231 | 0.23 | 0.563 |
| **Lead_Balloon_SummaryScore_DFE** | 0.576 (1.254) | -0.038 (0.658) | T(38)=1.939 | 0.06 | 1.330* |
| **2D_Manipulations_RT_DFE** | -0.006 (0.741) | 0.599 (1.892) | T(38)=-1.332 | 0.20 | 0.621 |
| **2D_Manipulations_SummaryScore_DFE** | 1.573 (1.484) | 1.236 (1.711) | T(38)=0.665 | 0.51 | 0.369 |
| **Motor_control_RT_DFE** | -0.097 (0.740) | -0.148 (0.766) | T(38)=0.214 | 0.83 | 0.314 |
| **Motor_Control_SummaryScore_DFE** | -0.012 (0.772) | -0.122 (0.764) | T(38)=0.453 | 0.65 | 0.335 |
| **Objects_memory_delayed_RT_DFE** | 0.416 (1.384) | -0.458 (0.898) | T(38)=2.369 | 0.02* | 2.653* |
| **Objects_memory_delayed_SummaryScore_DFE** | 0.073 (1.025) | -0.386 (1.866) | T(38)=0.964 | 0.34 | 0.446 |
| **Objects_memory_immediate_RT_DFE** | -0.106 (0.757) | -0.433 (0.840) | T(38)=1.293 | 0.20 | 0.596 |
| **Objects_memory_immediate_SummaryScore_DFE** | 0.023 (0.845) | -0.006 (1.268) | T(38)=0.085 | 0.93 | 0.310 |
| **Spotter_RT_DFE** | 0.360 (1.302) | -0.308 (0.841) | T(38)=1.927 | 0.06 | 1.303* |
| **Spotter_SummaryScore_DFE** | 0.093 (0.175) | 0.120 (0.361) | T(38)=-0.301 | 0.77 | 0.320 |
| **Verbal_analogies_RT_DFE** | 0.116 (0.743) | -0.027 (0.916) | T(38)=0.542 | 0.59 | 0.347 |
| **Verbal_analogies_SummaryScore_DFE** | -0.852 (1.355) | -0.592 (1.163) | T(38)=-0.651 | 0.52 | 0.366 |

**Supplementary Table S5:** **Associations between total extreme deviations in cortical thickness (positive and negative) and cognitive performance, fatigue, clinical symptom scores, cytokines, and routine blood biomarkers in the full sample.** Associations were quantified using Spearman’s rank correlations. Variables with p-values < 0.05 are highlighted in bold.

| **Variable** | **Stats** | **Total Extreme Positive Deviation Thickness** | **Total Extreme Negative Deviation Thickness** |
| --- | --- | --- | --- |
| Blocks_RT_DFE | Spearman's rho | 0.131 | 0.136 |
|  | p-value | 0.453 | 0.435 |
| BlocksSummaryScore_DFE | Spearman's rho | 0.022 | -0.103 |
|  | p-value | 0.899 | 0.558 |
| Lead_Balloon_SummaryScore_DFE | Spearman's rho | -0.127 | 0.179 |
|  | p-value | 0.468 | 0.304 |
| 2D_Manipulations_RT_DFE | Spearman's rho | -0.164 | 0.3 |
|  | p-value | 0.345 | 0.08 |
| 2D_Manipulations_SummaryScore_DFE | Spearman's rho | -0.139 | -0.332 |
|  | p-value | 0.425 | 0.051 |
| Motor_control_RT_DFE | Spearman's rho | -0.073 | -0.073 |
|  | p-value | 0.676 | 0.678 |
| Motor_Control_SummaryScore_DFE | Spearman's rho | -0.078 | -0.051 |
|  | p-value | 0.658 | 0.772 |
| Objects_memory_delayed_RT_DFE | Spearman's rho | 0.107 | 0.128 |
|  | p-value | 0.54 | 0.463 |
| Objects_memory_delayed_SummaryScore_DFE | Spearman's rho | -0.016 | -0.023 |
|  | p-value | 0.929 | 0.897 |
| Objects_memory_immediate_RT_DFE | Spearman's rho | -0.083 | -0.162 |
|  | p-value | 0.635 | 0.353 |
| Objects_memory_immediate_SummaryScore_DFE | Spearman's rho | 0.007 | -0.001 |
|  | p-value | 0.966 | 0.995 |
| Spotter_RT_DFE | Spearman's rho | 0.329 | 0.183 |
|  | p-value | 0.053 | 0.292 |
| Spotter_SummaryScore_DFE | Spearman's rho | **0.357** | -0.125 |
|  | p-value | **0.035** | 0.475 |
| Verbal_analogies_RT_DFE | Spearman's rho | 0.252 | -0.077 |
|  | p-value | 0.143 | 0.659 |
| Verbal_analogies_SummaryScore_DFE | Spearman's rho | -0.129 | -0.004 |
|  | p-value | 0.46 | 0.984 |
| IFNy | Spearman's rho | 0.223 | -0.015 |
|  | p-value | 0.228 | 0.935 |
| IL1b | Spearman's rho | 0.075 | -0.252 |
|  | p-value | 0.69 | 0.172 |
| IL6 | Spearman's rho | 0.211 | -0.128 |
|  | p-value | 0.247 | 0.484 |
| TNFa | Spearman's rho | 0.202 | -0.117 |
|  | p-value | 0.277 | 0.532 |
| IL10 | Spearman's rho | -0.01 | 0.088 |
|  | p-value | 0.958 | 0.643 |
| IL13 | Spearman's rho | 0.223 | -0.102 |
|  | p-value | 0.245 | 0.597 |
| IL8 | Spearman's rho | 0.04 | -0.307 |
|  | p-value | 0.832 | 0.094 |
| s100b | Spearman's rho | 0.202 | **0.346** |
|  | p-value | 0.26 | **0.049** |
| GFAP | Spearman's rho | -0.077 | 0.218 |
|  | p-value | 0.669 | 0.222 |
| Fatigue VAS | Spearman's rho | -0.19 | 0.015 |
|  | p-value | 0.275 | 0.933 |
| MOCA | Spearman's rho | -0.145 | -0.021 |
|  | p-value | 0.422 | 0.908 |
| PAIN detect score | Spearman's rho | -0.076 | 0.116 |
|  | p-value | 0.666 | 0.507 |
| WPI score | Spearman's rho | -0.218 | -0.176 |
|  | p-value | 0.209 | 0.312 |
| SS score | Spearman's rho | -0.067 | 0.234 |
|  | p-value | 0.703 | 0.175 |
| MFI_General fatigue | Spearman's rho | -0.114 | 0.106 |
|  | p-value | 0.521 | 0.552 |
| MFI_Physical fatigue | Spearman's rho | -0.093 | 0.03 |
|  | p-value | 0.599 | 0.867 |
| MFI_Reduced activity | Spearman's rho | -0.173 | -0.069 |
|  | p-value | 0.336 | 0.704 |
| MFI_Reduced motivation | Spearman's rho | -0.034 | -0.001 |
|  | p-value | 0.847 | 0.995 |
| MFI_Mental fatigue | Spearman's rho | **-0.487** | -0.148 |
|  | p-value | **0.003** | 0.403 |
| FAI_Total score | Spearman's rho | **-0.347** | -0.143 |
|  | p-value | **0.044** | 0.42 |
| HADS_A_Total | Spearman's rho | -0.252 | 0.018 |
|  | p-value | 0.15 | 0.921 |
| HADS_D_Total | Spearman's rho | -0.01 | 0.011 |
|  | p-value | 0.955 | 0.951 |
| ESS_total | Spearman's rho | -0.053 | 0.16 |
|  | p-value | 0.764 | 0.365 |
| PTSD_Total | Spearman's rho | -0.269 | 0.05 |
|  | p-value | 0.124 | 0.777 |
| DP_Unrefreshing_Sleep_Average_Severity | Spearman's rho | -0.072 | -0.046 |
|  | p-value | 0.685 | 0.798 |
| DP_PEM_Average_Severity | Spearman's rho | -0.217 | -0.119 |
|  | p-value | 0.217 | 0.504 |
| DP_Autonomic_Average_Severity | Spearman's rho | **-0.372** | 0.117 |
|  | p-value | **0.03** | 0.51 |
| DP_Neurocognitive_Average_Severity | Spearman's rho | -0.192 | 0.263 |
|  | p-value | 0.277 | 0.133 |
| DP_Immune_Average_Severity | Spearman's rho | **-0.4** | -0.029 |
|  | p-value | **0.019** | 0.871 |
| COMPASS_Total_Score | Spearman's rho | -0.17 | 0.209 |
|  | p-value | 0.338 | 0.235 |
| Global_PSQI_Score | Spearman's rho | -0.273 | 0.149 |
|  | p-value | 0.118 | 0.399 |
| Sodium | Spearman's rho | 0.208 | 0.008 |
|  | p-value | 0.262 | 0.967 |
| Creatinine | Spearman's rho | -0.076 | **0.449** |
|  | p-value | 0.683 | **0.011** |
| Proteins_total | Spearman's rho | 0.1 | -0.189 |
|  | p-value | 0.591 | 0.31 |
| Albumin | Spearman's rho | 0.258 | -0.141 |
|  | p-value | 0.162 | 0.449 |
| Bilirubin_total | Spearman's rho | -0.312 | -0.031 |
|  | p-value | 0.107 | 0.876 |
| ALP | Spearman's rho | 0.054 | -0.081 |
|  | p-value | 0.773 | 0.666 |
| AST | Spearman's rho | **-0.367** | -0.29 |
|  | p-value | **0.042** | 0.114 |
| Gamma_GT | Spearman's rho | -0.279 | -0.043 |
|  | p-value | 0.128 | 0.818 |
| CRP_recoded | Spearman's rho | -0.136 | 0.313 |
|  | p-value | 0.466 | 0.086 |
| Globulin | Spearman's rho | -0.149 | -0.142 |
|  | p-value | 0.423 | 0.447 |
| Ferritin | Spearman's rho | -0.195 | 0.255 |
|  | p-value | 0.294 | 0.167 |
| Hemoglobin | Spearman's rho | -0.012 | -0.075 |
|  | p-value | 0.951 | 0.687 |
| MCHC | Spearman's rho | 0.147 | -0.246 |
|  | p-value | 0.43 | 0.182 |
| Platelet_count | Spearman's rho | -0.005 | 0.102 |
|  | p-value | 0.979 | 0.587 |
| ESR | Spearman's rho | -0.265 | -0.237 |
|  | p-value | 0.157 | 0.208 |
| Time from 1st infection (months) | Spearman's rho | -0.091 | -0.122 |
|  | p-value | 0.604 | 0.485 |
| Potassium | Spearman's rho | 0.266 | 0.329 |
|  | p-value | 0.164 | 0.081 |
| Urea | Spearman's rho | -0.06 | **0.367** |
|  | p-value | 0.747 | **0.042** |
| Glucose | Spearman's rho | -0.077 | 0.083 |
|  | p-value | 0.682 | 0.655 |
| Calcium_corrected | Spearman's rho | 0.066 | -0.182 |
|  | p-value | 0.724 | 0.328 |
| Phosphatase | Spearman's rho | 0.167 | 0.025 |
|  | p-value | 0.369 | 0.895 |
| Magnesium | Spearman's rho | **0.368** | 0.258 |
|  | p-value | **0.042** | 0.161 |
| WCC | Spearman's rho | 0.231 | -0.169 |
|  | p-value | 0.211 | 0.363 |
| RCC | Spearman's rho | -0.149 | -0.083 |
|  | p-value | 0.425 | 0.655 |
| Haematocrit | Spearman's rho | -0.125 | -0.037 |
|  | p-value | 0.503 | 0.845 |
| MCV | Spearman's rho | 0.162 | 0.142 |
|  | p-value | 0.383 | 0.448 |
| MCH | Spearman's rho | 0.242 | -0.027 |
|  | p-value | 0.191 | 0.883 |
| RDW | Spearman's rho | 0.185 | 0.063 |
|  | p-value | 0.32 | 0.736 |
| MPV | Spearman's rho | 0.163 | -0.33 |
|  | p-value | 0.381 | 0.07 |
| Neutrophils | Spearman's rho | 0.278 | -0.164 |
|  | p-value | 0.13 | 0.377 |
| Lymphocytes | Spearman's rho | 0.1 | -0.143 |
|  | p-value | 0.592 | 0.442 |
| Monocytes | Spearman's rho | 0.298 | -0.063 |
|  | p-value | 0.104 | 0.737 |
| Eosinophils | Spearman's rho | 0.146 | **0.418** |
|  | p-value | 0.435 | **0.019** |
| Basophils | Spearman's rho | 0.242 | 0.13 |
|  | p-value | 0.189 | 0.486 |
| Immature_Granulocyte_Count | Spearman's rho | -0.094 | -0.079 |
|  | p-value | 0.616 | 0.673 |
| TSH | Spearman's rho | 0.11 | 0.272 |
|  | p-value | 0.556 | 0.139 |
| T4_free | Spearman's rho | -0.066 | -0.111 |
|  | p-value | 0.726 | 0.554 |
| NLR | Spearman's rho | 0.186 | -0.071 |
|  | p-value | 0.317 | 0.703 |
| MLR | Spearman's rho | 0.142 | 0.094 |
|  | p-value | 0.445 | 0.616 |

***Supplementary Figure S1.* Regional differences in cortical thickness, surface area, and subcortical volumes deviation scores between post-COVID-19 patients and healthy controls**. Comparisons were adjusted for age, gender, handedness, total intracranial volume (TIV) and the presence of anosmia/ageusia. Colours indicate effect sizes (Cohen’s d), with red representing increased measures and blue indicating decreased measures in post-COVID-19 patients compared to controls. Significant results are thresholded at uncorrected p < 0.05.

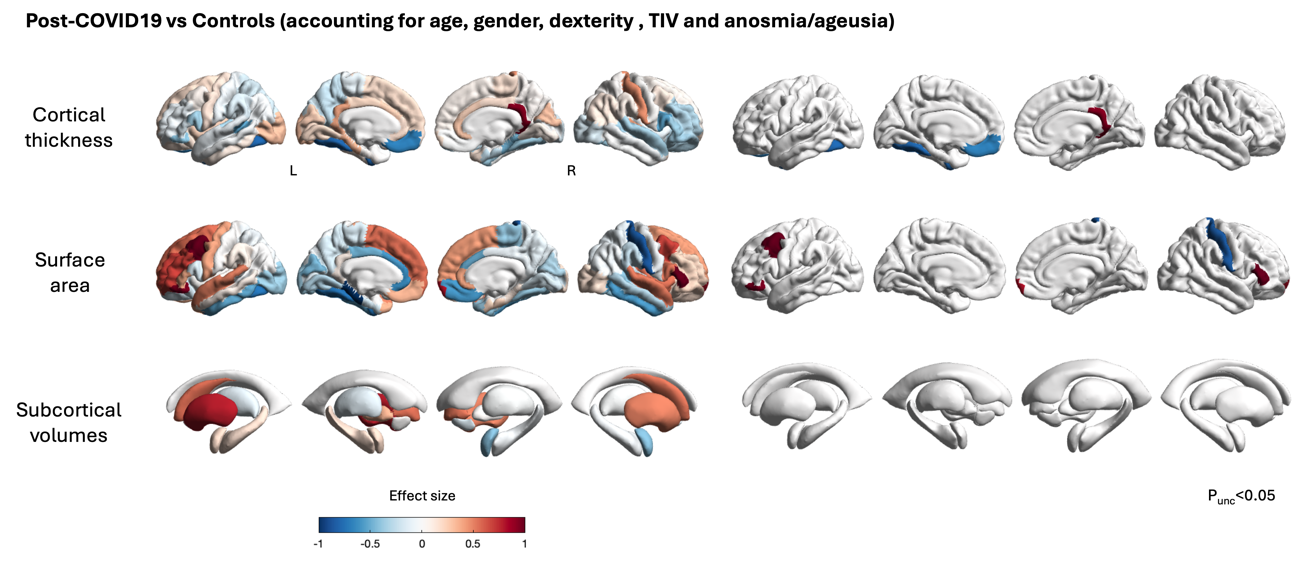

**Supplementary Table S6. Regional group differences in cortical thickness and surface area between PCS and control participants, analysed with and without accounting for self-reported anosmia/ageusia as a covariate.** Effect sizes (Cohen’s d) and uncorrected p-values are reported for each cortical and subcortical region, based on general linear models adjusting for age, sex, handedness, and total intracranial volume (TIV).

| **Region** | **Without accounting for anosmia/ageusia** | | **Accounting for anosmia/ageusia** | |
| --- | --- | --- | --- | --- |
|  | **Effect Size** | **p-value** | **Effect Size** | **Group p-value** |
| L_bankssts_thickavg | -0.208200066 | 0.571540362 | -0.524055826 | 0.276027092 |
| L_caudalanteriorcingulate_thickavg | 0.407518609 | 0.217333414 | 0.245632139 | 0.563773455 |
| L_caudalmiddlefrontal_thickavg | -0.057622993 | 0.815080424 | 0.07004802 | 0.829361686 |
| L_cuneus_thickavg | -0.015351183 | 0.963927025 | 0.234765823 | 0.600422736 |
| L_entorhil_thickavg | 0.148940856 | 0.530541929 | 0.043263295 | 0.883171652 |
| L_fusiform_thickavg | -0.568890681 | 0.020762448 | -0.795345774 | 0.013732572 |
| L_inferiorparietal_thickavg | 0.126317956 | 0.610111027 | -0.038735252 | 0.905399148 |
| L_inferiortemporal_thickavg | 0.109341321 | 0.565341715 | 0.112104371 | 0.659318253 |
| L_isthmuscingulate_thickavg | 0.306269578 | 0.257046997 | 0.375725003 | 0.293244666 |
| L_lateraloccipital_thickavg | 0.509273922 | 0.028106772 | 0.300773156 | 0.291813608 |
| L_lateralorbitofrontal_thickavg | 0.067966758 | 0.803757485 | 0.136767359 | 0.684883505 |
| L_lingual_thickavg | 0.141346044 | 0.613238568 | 0.177878014 | 0.633399391 |
| L_medialorbitofrontal_thickavg | -0.602780145 | 0.006627927 | -0.669418408 | 0.021954385 |
| L_middletemporal_thickavg | -0.175382718 | 0.488879602 | -0.024002482 | 0.942436362 |
| L_parahippocampal_thickavg | 0.396462155 | 0.096199122 | 0.333775514 | 0.288543831 |
| L_paracentral_thickavg | -0.050605374 | 0.872049334 | -0.105384297 | 0.801936833 |
| L_parsopercularis_thickavg | -0.350133099 | 0.199053279 | -0.353262817 | 0.298385642 |
| L_parsorbitalis_thickavg | -0.451183686 | 0.101085439 | -0.410526364 | 0.259617771 |
| L_parstriangularis_thickavg | -0.25217934 | 0.228547147 | -0.141186633 | 0.601958495 |
| L_pericalcarine_thickavg | 0.348361103 | 0.336010602 | 0.18626915 | 0.69571967 |
| L_postcentral_thickavg | 0.066479449 | 0.722240914 | -0.065965974 | 0.789747 |
| L_posteriorcingulate_thickavg | 0.403464952 | 0.206218266 | 0.202909456 | 0.628378709 |
| L_precentral_thickavg | -0.176544755 | 0.595450009 | 0.155794924 | 0.709835759 |
| L_precuneus_thickavg | -0.100449034 | 0.690957937 | -0.25678481 | 0.429671334 |
| L_rostralanteriorcingulate_thickavg | 0.071512117 | 0.7788415 | 0.165449463 | 0.626616771 |
| L_rostralmiddlefrontal_thickavg | -0.092245738 | 0.685402995 | -0.078752684 | 0.794147111 |
| L_superiorfrontal_thickavg | -0.003617638 | 0.99117024 | 0.178232738 | 0.679613848 |
| L_superiorparietal_thickavg | -0.108014647 | 0.626920103 | -0.202203419 | 0.495077243 |
| L_superiortemporal_thickavg | 0.297453734 | 0.266997364 | -0.066680016 | 0.844789303 |
| L_supramargil_thickavg | -0.059428017 | 0.807439018 | -0.160824685 | 0.620488073 |
| L_frontalpole_thickavg | 0.090848391 | 0.7632465 | 0.128009322 | 0.743777981 |
| L_temporalpole_thickavg | 0.414373562 | 0.230915227 | 0.004564518 | 0.991593726 |
| L_transversetemporal_thickavg | -0.114972492 | 0.673714287 | -0.338067276 | 0.338889317 |
| L_insula_thickavg | 0.069046148 | 0.819956654 | 0.222921274 | 0.581658471 |
| R_bankssts_thickavg | -0.052882523 | 0.861105279 | -0.349794613 | 0.353074408 |
| R_caudalanteriorcingulate_thickavg | 0.174543277 | 0.433701462 | 0.225758341 | 0.445901829 |
| R_caudalmiddlefrontal_thickavg | -0.327599459 | 0.320719931 | 0.12573487 | 0.76438595 |
| R_cuneus_thickavg | 0.404139684 | 0.233203334 | 0.292895958 | 0.51422735 |
| R_entorhil_thickavg | -0.117707888 | 0.69980038 | -0.247922565 | 0.531066408 |
| R_fusiform_thickavg | -0.383746542 | 0.135179968 | -0.361120685 | 0.276956881 |
| R_inferiorparietal_thickavg | 0.092185938 | 0.660435798 | 0.125550256 | 0.653157855 |
| R_inferiortemporal_thickavg | 0.224169023 | 0.36188204 | -0.002764423 | 0.993105864 |
| R_isthmuscingulate_thickavg | 0.524862007 | 0.057986819 | 0.933574615 | 0.008629638 |
| R_lateraloccipital_thickavg | 0.218783062 | 0.426665653 | -0.169928216 | 0.624737256 |
| R_lateralorbitofrontal_thickavg | -0.387190997 | 0.041538276 | -0.304058061 | 0.202419193 |
| R_lingual_thickavg | 0.270239042 | 0.418231141 | 0.09348065 | 0.828335231 |
| R_medialorbitofrontal_thickavg | -0.050318602 | 0.823103644 | -0.00422664 | 0.988714038 |
| R_middletemporal_thickavg | 0.047983727 | 0.863449986 | -0.036891566 | 0.920626387 |
| R_parahippocampal_thickavg | -0.128020901 | 0.573218526 | -0.464282065 | 0.110551824 |
| R_paracentral_thickavg | -0.060260355 | 0.84673888 | 0.164052537 | 0.688269895 |
| R_parsopercularis_thickavg | -0.724379874 | 0.023437572 | -0.527254137 | 0.188532702 |
| R_parsorbitalis_thickavg | -0.742451902 | 0.001043271 | -0.349356821 | 0.168873096 |
| R_parstriangularis_thickavg | -0.320040471 | 0.229335234 | -0.363521451 | 0.3062155 |
| R_pericalcarine_thickavg | 0.340716055 | 0.39164396 | -0.026651078 | 0.958872961 |
| R_postcentral_thickavg | 0.57372741 | 0.046208571 | 0.449094732 | 0.230627496 |
| R_posteriorcingulate_thickavg | 0.121854898 | 0.615297538 | 0.169313086 | 0.600731852 |
| R_precentral_thickavg | -0.521348942 | 0.189369479 | 0.02149348 | 0.964969078 |
| R_precuneus_thickavg | 0.202204605 | 0.423918246 | 0.045156232 | 0.892124438 |
| R_rostralanteriorcingulate_thickavg | 0.321356058 | 0.128983862 | 0.275756201 | 0.325265601 |
| R_rostralmiddlefrontal_thickavg | -0.247429693 | 0.169920711 | -0.303461632 | 0.192964612 |
| R_superiorfrontal_thickavg | -0.410684467 | 0.132207673 | 0.067504164 | 0.839100949 |
| R_superiorparietal_thickavg | -0.036498341 | 0.870504764 | 0.024503414 | 0.934260702 |
| R_superiortemporal_thickavg | -0.095167892 | 0.677183855 | -0.329596159 | 0.27290028 |
| R_supramargil_thickavg | 0.162435937 | 0.448513202 | 0.088046234 | 0.753955766 |
| R_frontalpole_thickavg | -0.134838894 | 0.617054874 | 0.161273179 | 0.641297009 |
| R_temporalpole_thickavg | 0.542171255 | 0.132460937 | 0.141801315 | 0.753180667 |
| R_transversetemporal_thickavg | 0.121840233 | 0.713809894 | 0.403743013 | 0.356131181 |
| R_insula_thickavg | 0.035316046 | 0.907778386 | -0.090483581 | 0.82084964 |
| L_bankssts_surfavg | 0.09904335 | 0.759281342 | -0.052025401 | 0.898687988 |
| L_caudalanteriorcingulate_surfavg | -0.377660189 | 0.294784709 | -0.592340741 | 0.208703023 |
| L_caudalmiddlefrontal_surfavg | 0.157521563 | 0.664882618 | 1.239528789 | 0.002321619 |
| L_cuneus_surfavg | -0.206939515 | 0.55641489 | -0.454660623 | 0.32564649 |
| L_entorhil_surfavg | 0.258461961 | 0.543621427 | -0.183529977 | 0.740835416 |
| L_fusiform_surfavg | 0.041774738 | 0.902244025 | -0.732549511 | 0.070304074 |
| L_inferiorparietal_surfavg | 0.006316202 | 0.984926679 | 0.000602215 | 0.998917322 |
| L_inferiortemporal_surfavg | -0.244671651 | 0.449880253 | -0.517590989 | 0.226632727 |
| L_isthmuscingulate_surfavg | -0.148171157 | 0.692454978 | 0.095771369 | 0.846854716 |
| L_lateraloccipital_surfavg | -0.048790271 | 0.879817602 | -0.363820752 | 0.390306058 |
| L_lateralorbitofrontal_surfavg | -0.349039903 | 0.23343447 | -0.175981285 | 0.648110272 |
| L_lingual_surfavg | -0.197168178 | 0.549822337 | -0.291422462 | 0.508568936 |
| L_medialorbitofrontal_surfavg | 0.237679802 | 0.374694798 | 0.340802446 | 0.340647745 |
| L_middletemporal_surfavg | 0.116188929 | 0.702059225 | 0.091367327 | 0.821310531 |
| L_parahippocampal_surfavg | -0.592088627 | 0.167053133 | -1.019517051 | 0.069358902 |
| L_paracentral_surfavg | -0.087100735 | 0.770888474 | -0.167211631 | 0.670800939 |
| L_parsopercularis_surfavg | -0.072750731 | 0.846302636 | -0.162307505 | 0.742662436 |
| L_parsorbitalis_surfavg | 0.344408523 | 0.351585064 | 0.967637001 | 0.038611726 |
| L_parstriangularis_surfavg | 0.099614609 | 0.747016503 | 0.361758897 | 0.375078776 |
| L_pericalcarine_surfavg | -0.181440597 | 0.617242415 | -0.141328585 | 0.769688424 |
| L_postcentral_surfavg | 0.122364796 | 0.643751515 | 0.012118953 | 0.972440472 |
| L_posteriorcingulate_surfavg | -0.582054573 | 0.117607813 | -0.479114546 | 0.329093767 |
| L_precentral_surfavg | 0.112132442 | 0.690516147 | 0.395409438 | 0.283215258 |
| L_precuneus_surfavg | -0.014162013 | 0.953586768 | -0.184279422 | 0.566936827 |
| L_rostralanteriorcingulate_surfavg | -0.302986253 | 0.323222166 | -0.61941355 | 0.126180323 |
| L_rostralmiddlefrontal_surfavg | 0.179596999 | 0.624923652 | 0.680553572 | 0.152577738 |
| L_superiorfrontal_surfavg | 0.352504459 | 0.358461134 | 0.551407371 | 0.277528398 |
| L_superiorparietal_surfavg | 0.006147727 | 0.984292504 | -0.067872106 | 0.870882183 |
| L_superiortemporal_surfavg | 0.257463659 | 0.33272084 | 0.493887019 | 0.148851435 |
| L_supramargil_surfavg | -0.074856976 | 0.828471712 | 0.098838943 | 0.829744223 |
| L_frontalpole_surfavg | -0.253576358 | 0.496467289 | 0.557005699 | 0.214051067 |
| L_temporalpole_surfavg | 0.143440943 | 0.687821264 | 0.344934149 | 0.4678363 |
| L_transversetemporal_surfavg | -0.06363861 | 0.841721265 | 0.058733481 | 0.884469704 |
| L_insula_surfavg | -0.143552751 | 0.587086193 | 0.016072022 | 0.963293056 |
| R_bankssts_surfavg | 0.048687769 | 0.894451889 | -0.409055184 | 0.384159276 |
| R_caudalanteriorcingulate_surfavg | -0.416703464 | 0.261889289 | -0.499626622 | 0.312731016 |
| R_caudalmiddlefrontal_surfavg | 0.505536245 | 0.152642121 | 0.674305327 | 0.142523781 |
| R_cuneus_surfavg | -0.338941913 | 0.314081193 | -0.319159528 | 0.475082535 |
| R_entorhil_surfavg | 0.19894598 | 0.66337892 | 0.213831295 | 0.725971633 |
| R_fusiform_surfavg | -0.201362111 | 0.570719704 | -0.379012434 | 0.419074948 |
| R_inferiorparietal_surfavg | -0.026273463 | 0.936939939 | -0.061623969 | 0.888352748 |
| R_inferiortemporal_surfavg | -0.260213374 | 0.498896469 | -0.571628358 | 0.249863616 |
| R_isthmuscingulate_surfavg | -0.000850682 | 0.997514878 | -0.071940264 | 0.843111373 |
| R_lateraloccipital_surfavg | 0.497343593 | 0.198630773 | 0.15739146 | 0.753243978 |
| R_lateralorbitofrontal_surfavg | -0.070182992 | 0.831229227 | 0.171781645 | 0.693342531 |
| R_lingual_surfavg | 0.029007824 | 0.940758615 | -0.168645942 | 0.734171088 |
| R_medialorbitofrontal_surfavg | -0.225610136 | 0.498997904 | -0.590364007 | 0.179048601 |
| R_middletemporal_surfavg | 0.174871104 | 0.624788021 | -0.291110649 | 0.502950833 |
| R_parahippocampal_surfavg | -0.26843003 | 0.287095582 | 0.126852945 | 0.688165435 |
| R_paracentral_surfavg | -0.252099178 | 0.260703076 | -0.382992181 | 0.199878348 |
| R_parsopercularis_surfavg | 0.637822854 | 0.104143771 | 0.531247466 | 0.304811317 |
| R_parsorbitalis_surfavg | 0.062802744 | 0.815372425 | 0.210446186 | 0.55677834 |
| R_parstriangularis_surfavg | 0.414339064 | 0.233096026 | 1.031509211 | 0.019943789 |
| R_pericalcarine_surfavg | -0.269817736 | 0.477085875 | -0.208167072 | 0.676071618 |
| R_postcentral_surfavg | -0.547963348 | 0.045975361 | -0.842322607 | 0.019542634 |
| R_posteriorcingulate_surfavg | 0.004687441 | 0.989207099 | -0.0826453 | 0.858070691 |
| R_precentral_surfavg | 0.23597657 | 0.545784445 | -0.048292763 | 0.925031649 |
| R_precuneus_surfavg | -0.256419626 | 0.431590263 | -0.105656076 | 0.799555677 |
| R_rostralanteriorcingulate_surfavg | 0.214163463 | 0.49917905 | 0.38287129 | 0.365089663 |
| R_rostralmiddlefrontal_surfavg | -0.153596434 | 0.688727827 | 0.209358953 | 0.676089507 |
| R_superiorfrontal_surfavg | 0.543811368 | 0.191890966 | 0.419913787 | 0.444367756 |
| R_superiorparietal_surfavg | -0.327125347 | 0.289357995 | -0.09031583 | 0.819487071 |
| R_superiortemporal_surfavg | 0.345469316 | 0.41624684 | 0.521900352 | 0.345445676 |
| R_supramargil_surfavg | -0.150219536 | 0.648485958 | -0.236622305 | 0.590474189 |
| R_frontalpole_surfavg | 0.626448667 | 0.016390808 | 0.807354635 | 0.01939865 |
| R_temporalpole_surfavg | -0.46704481 | 0.235143738 | -0.37197924 | 0.471148784 |
| R_transversetemporal_surfavg | -0.353349818 | 0.287318236 | -0.200585366 | 0.647379951 |
| R_insula_surfavg | 0.213557527 | 0.502331074 | 0.541649189 | 0.197303427 |
| Laccumb | 0.024735506 | 0.946094696 | 0.009440321 | 0.984591582 |
| Lamyg | 0.028951963 | 0.919896721 | 0.179687696 | 0.637291428 |
| Lcaud | 0.242705292 | 0.450572135 | 0.547669121 | 0.195685671 |
| Lhippo | 0.333456312 | 0.234970567 | 0.165297989 | 0.652409515 |
| Lpal | 0.161110376 | 0.546267961 | 0.34149353 | 0.331892501 |
| Lput | 0.552442012 | 0.079771752 | 0.774681771 | 0.064158531 |
| Lthal | 0.074131452 | 0.784002984 | -0.116299063 | 0.732655579 |
| Raccumb | 0.206364553 | 0.511735385 | -0.136930743 | 0.737084308 |
| Ramyg | -0.198317074 | 0.467349191 | -0.427481983 | 0.236481924 |
| Rcaud | 0.231992892 | 0.458136663 | 0.53177834 | 0.196028808 |
| Rhippo | 0.171533899 | 0.529822918 | -0.035570829 | 0.919357375 |
| Rpal | -0.045610232 | 0.891227323 | 0.025110095 | 0.953454865 |
| Rput | 0.180667146 | 0.648735296 | 0.495107132 | 0.346975655 |
| Rthal | -0.118603847 | 0.637275009 | -0.00977145 | 0.976177029 |
